# Redox distribution of Asgard archaea and co-occurring taxa in microbial mats from an early Proterozoic ecosystem analog

**DOI:** 10.64898/2026.03.20.713109

**Authors:** Ana Gutiérrez-Preciado, Agathe Struillou, Lewen Liang, Miguel Iniesto, Philippe Deschamps, Laura Eme, Yvan Zivanovic, José M. López-García, Karim Benzerara, David Moreira, Purificación López-García

## Abstract

Increasing evidence supports that eukaryotes originated from the symbiosis of an Asgard archaeon, the alphaproteobacterial ancestor of mitochondria, and possibly additional bacterial contributions. Eukaryogenesis likely occurred in redox-transition environments such as microbial mats or shallow sediments ∼2 billion years ago, when atmospheric oxygen was far lower than today, and oceans were euxinic. We investigated Asgardarchaeota-enriched microbial mats from the low-oxygen, sulfidic Catherine volcano lake (Afar region, Ethiopia), mimicking early Proterozoic conditions. 16S rRNA gene metabarcoding, metagenomics, and metagenome-assembled genome (MAG) analyses across redox-stratified layers of in situ and mesocosm-maintained mats revealed that Asgardarchaeota thrived in the sulfate-reduction zone, mainly co-occurring with Desulfurobacterota and Myxococcota, among others. Lokiarchaeia and Thorarchaeia were more abundant in deeper, anoxic layers. Within Heimdallarchaeia, Heimdallarchaeales were enriched in upper layers, correlating with oxygen-tolerant hydrogenase and sulfate-reduction genes, whereas Hodarchaeales were more relatively abundant in anoxic layers, correlating with methanogenesis. Although reactive-oxygen-species defense mechanisms were widespread in Asgardarchaeota MAGs, they lacked genes for aerobic respiration. These results support the hypothesis that Asgard archaea engaged primarily in syntrophic interactions with sulfate-reducers under conditions prevailing on the early Earth at the onset of eukaryogenesis.

---

The discovery of Asgardarchaeota (informally, Asgard archaea) shed new light on the origin of the eukaryotic cell^1–5^. Known for the past two decades from their 16S rRNA gene sequences^6–8^, Asgard archaea gained widespread attention about a decade ago with the reconstruction of the first metagenome-assembled genomes (MAGs) for the Lokiarchaeia, the first identified Asgardarchaeota group, from deep-sea sediment metagenomes^9^. This and subsequent similar studies^10–14^ expanded the known diversity for the clade and revealed that Asgardarchaeota genomes encode highly similar homologs to eukaryotic proteins, in numbers exceeding those shared between eukaryotes and other archaeal lineages. Furthermore, phylogenomic analyses of conserved informational proteins place eukaryotes within Asgardarchaeota, either as sister to the class Heimdallarchaeia^15^, or to the heimdallarcheial order Hodarchaeales^16^. These observations support the notion that eukaryotes evolved by the symbiotic merging of one Asgard archaeon and at least the alphaproteobacterial ancestor of mitochondria^2,9^, possibly via some form of metabolic symbiosis or syntrophy^17,18^. While most eukaryogenesis models involve only these two symbiotic partners, others invoke additional permanent or transient symbioses with one sulfate-reducing bacterium and/or other bacteria^19–21^. Ancestral gene content inference indeed suggests a more or less substantial contribution of non-alphaproteobacterial genes to the Last Eukaryotic Common Ancestor (LECA) in addition to asgardarchaeal genes^22–25^. Whether those genes were acquired from bacterial symbiotic partners, by random horizontal gene transfer from bacteria co-existing in the same environment^26^, or a combination of both, is an open question that a better knowledge of the ecology of Asgardarchaeotaand their co-occurring taxa could help answer.

The known diversity of Asgard archaea has continuously increased in recent years, as well as their environmental distribution. Currently, the phylum Asgardarchaeota comprises 12 classes according to the Genome Taxonomy Database (GTDB, r226)^27^. Most asgardarchaeal MAGs have been assembled from oxygen-deprived environments, including deep-sea and coastal marine sediments, lake sediments, hot springs and microbial mats^10,28,29^. However, Heimdallarchaeia have also been detected in microoxic environments^30,31^. Diverse Heimdallarchaeia encode proteins involved in the detoxification of oxygen reactive species (ROS), haem biosynthesis and an electron transport chain complex (IV) suggestive of aerobic respiration potential^31^. While aerobic respiration remains to be experimentally demonstrated, some Heimdallarchaiea are aerotolerant, notably in the presence of aerobic microorganisms^32^, and the possession of terminal oxidases and globins indicates oxygen-adaptive plasticity^33^. Many Asgardarchaeotaappear to prefer slightly saline environments, ranging from brackish and marine sediments^10–12,34^ to sediments^35,36^ and microbial mats^37–39^ in lagoons, saline lakes and ponds.

From a metabolic perspective, Asgardarchaeota are inferred to be mostly heterotrophic, intervening in the anaerobic degradation of organic matter^11^. Some members might also fix carbon and be mixotrophic and/or chemoautotrophic^13,40^; many carry light-driven rhodopsins^41^. Asgardarchaeota seem to be either hydrogen/electron donors or acceptors, being likely involved in interspecies hydrogen syntrophies under their natural anoxic conditions^13,17^. Confirming this prediction, the first Asgardarchaeota enriched in culture, including Lokiarchaeia and Heimdallarchaeia (Hodarchaeales) members, grow in syntrophy with sulfate-reducing bacteria (Desulfobacterota), methanogenic archaea or both^19,42–44^. Nonetheless, enrichment cultures generally comprise antibiotic treatment, which might affect original interactions, and information about potential symbiotic partners of Asgardarchaeota in natural ecosystems is still missing.

Eukaryotes likely evolved during the early Proterozoic, little after the Great Oxidation Event (GOE, ∼2.4 Ga) that resulted in the oxygenation of the atmosphere^45–47^. However, atmospheric oxygen levels remained relatively low, and the deep ocean anoxic, until the Neoproterozoic (∼0.8-0.5 Ga). Thus, at the time when eukaryogenesis started, surface waters on the planet were suboxic, having at most ∼10% of today’s atmospheric oxygen saturation and undergoing pulses of anoxia^46^. At the same time, early continental oxidative weathering made the late Archaean-early Proterozoic oceans euxinic^48,49^ and, possibly, ∼1.5-2 times more saline than modern oceans^50^. Therefore, eukaryotes likely evolved in sulfidic, moderately salty and relatively oxygen-poor environments, such as microbial mats or shallow sediments, where Asgardarchaeota could interact with the facultatively aerobic ancestor of mitochondria^21,31,46^. Here, we study Asgard archaea–enriched microbial mats from the suboxic and geothermally active crater lake (DAN-LK4) on Catherine volcano, south of the Danakil Depression, Ethiopia^51^. Located at the north of the Erta Ale volcanic chain, south to the hypersaline Lake Karum and the polyextreme Dallol geothermal dome^52,53^, this sulfidic, saline and oxygen-poor ecosystem represents a suitable analog of early Proterozoic ecosystems. Using 16S rRNA gene metabarcoding and genome-resolved metagenomics, we characterize the microbial community composition and functional potential of different mat layers along the vertical redox gradient in natural and mesocosm-maintained samples. We show that, at phylum level, community structure is relatively preserved in deep anoxic layers, but exhibits differences in surface layers and at the level of lower-ranking taxa. Co-occurrence networks from metabarcoding and metagenomic data of cultured and in situ microbial mat layers reveal shared ecological redox preferences of Asgardarchaeota and neighboring taxa and identify candidate taxa with which Asgardarchaeota might engage in potential symbiotic interactions.

## Results

### Microbial mats from an early Proterozoic ecosystem analog

The Catherine and Gada Ale volcanos lie at the north end of the Erta Ale volcanic chain, immediately to the south of the Danakil Depression salt desert, a subaerial tectonically controlled basin at a depth of 120 m below sea level (mbsl), and the saline-to-hypersaline lakes Bakili and Karum^52,54^ (Fig.1a). This arid area of the East African Rift, located at the confluence of three tectonic plates (the Afar triangle), undergoes active rifting and volcanism^54,55^. The Catherine volcano has a characteristic circular low and wide cone of gentle external slopes, 500 m in diameter at the top, 1 km at the base, peaking 60 m over the basaltic ground (70 m above the bottom of the Danakil Depression, i.e. 50 mbsl). Its morphology and composition (basaltic glassy fragments or hyaloclastites) is characteristic of tuff rings resulting from magma interaction with surface water^56^, testifying to its submarine formation during ancient marine intrusions into the Danakil Depression. The Catherine volcano is surrounded by more recent subaerial lavas of the Gada Ale^54^. While the Gada Ale’s crater hosts a boiling mud lake, with sulfur as main sublimate, the Catherine volcano crater harbors a water lake ∼65 m below the crater rim, (i.e. 115 mbsl) fed by subaquatic hot springs^54^ (Fig.1a). At the time of sampling, the lake was continuously bubbling and strongly smelled of H_2_S (<u>Supplementary Video 1</u>). Physicochemical parameters and hydrochemical analyses revealed a neutral (pH 7.3), saline (104.5 ppt) lake enriched in Na-Ca-Mg-K salts and rare elements (W, Rb, Ba, among others) (Supplementary Table 1). Surface waters were poorly oxygenated (0.8 mg/l, i.e. ∼10-15% of normally oxygenated waters) and displayed low redox potential (10.7 mV, i.e. ∼2.5% of seawater). These conditions are reminiscent of those prevailing in late Archaean-early Proterozoic aquatic systems.

The bottom of the lake, at least close to the shore, was covered by a thick, consistent microbial mat displaying dark layers (Fig.1a; <u>Supplementary Video 1</u>). We collected a relatively large fragment of this mat (∼7×18×25 cm), which was transported back to the laboratory and used to set a small mesocosm. Although environmental conditions changed from those in the natural setting, incubation conditions were stable and mimicked to some extent the original ones: immersion in DAN-LK4 water (mat covered by ∼3-5 mm water, as at the collection point in the field), circumneutral pH, 12-h daylight period, and ∼25°CWe subsampled layers of the microbial mat in situ (DAN-LK4m1 to DAN-LK4m4; fixed at the time of collection) and equivalent layers (DAN-LK4m1cu to DAN-LK4m4cu) after 35 months of incubation in the culture room (Fig. 1a) for subsequent metabarcoding and metagenomic analyses. To determine the redox potential across mat layers, we measured pH, oxygen and sulfide concentrations in triplicate using microelectrodes in the mesocosm-maintained microbial mat. While pH remained stable at ∼6, O_2_ decreased rapidly to undetectable levels below 1 cm, while H_2_S increased to broadly peak between 2-4 cm, indicating a sulfate-reduction zone maximum (Fig.1b).

**Fig. 1.**
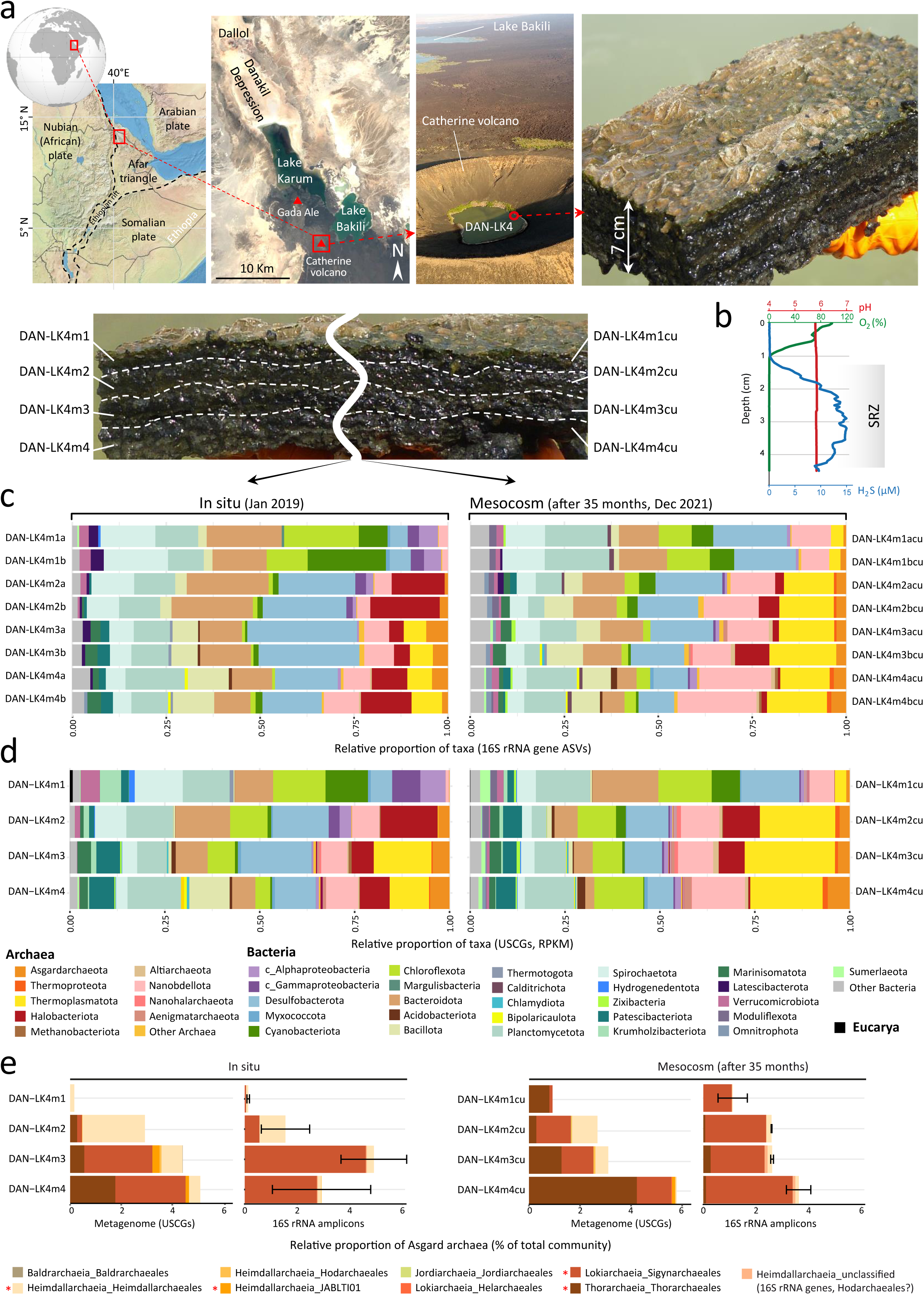
Sampling location and microbial mat community structure in the Catherine volcano lake (DAN-LK4) inferred from metabarcoding and metagenomic data of mat layers as collected in situ and after 35 months of mesocosm incubation. a, Sampling site at the Catherine volcano at the northern tip of the Erta Ale volcanic chain, southwest of Lake Bakili and aligned with the Danakil Depression, Ethiopia. The two leftmost panels were created using images from Wikimedia Commons and Google Earth, respectively. Photographs taken on the sampling day, showing the Catherine volcano and the studied microbial mat with its stratified layers, are displayed on the right and bottom panels. b, Vertical profile of pH, O_2_ and H_2_S levels in the mesocosm microbial mat measured with microelectrodes. SRZ, sulfate-reduction zone. c, Barplots showing the prokaryotic community composition of microbial mat layers (in replicates, labeled a and b) of in situ (left) and mesocosm-incubated (right) microbial mats based on 16S rRNA gene amplicon metabarcoding. ASV, amplicon sequence variant. d, Barplots showing the microbial community composition (archaea, bacteria, eukaryotes) based on the proportion of universal single-copy genes (USCGs; ribosomal proteins), expressed in reads per kilobase gene length per million reads (RPKMs), in metagenomes of the different microbial mat layers. The taxonomic affiliation of ASVs and USCGs in B and C is given at the phylum level except for the classes (c_) Alphaproteobacteria and Gammaproteobacteria (phylum Pseudomonadota) and eukaryotes, grouped as a single domain, Eucarya. Taxonomy follows the GTDB r226 nomenclature. Only phyla in proportions higher than 0.5% (16S rRNA gene-based ASVs or USCGs) are given, the rest are classed as ‘other bacteria’ or ‘other archaea’. e, Barplots showing the relative proportion of Asgardarchaeota at the order level as inferred from metagenome data (% USCGs, RPKMs) and 16S rRNA metabarcoding data (% ASVs). Orders detected by both approaches are labeled with an asterisk. The category Heimdallarchaeia_unclassified identified by 16S rRNA gene sequencing likely corresponds to Hodarchaeales.

**Fig. 2.**
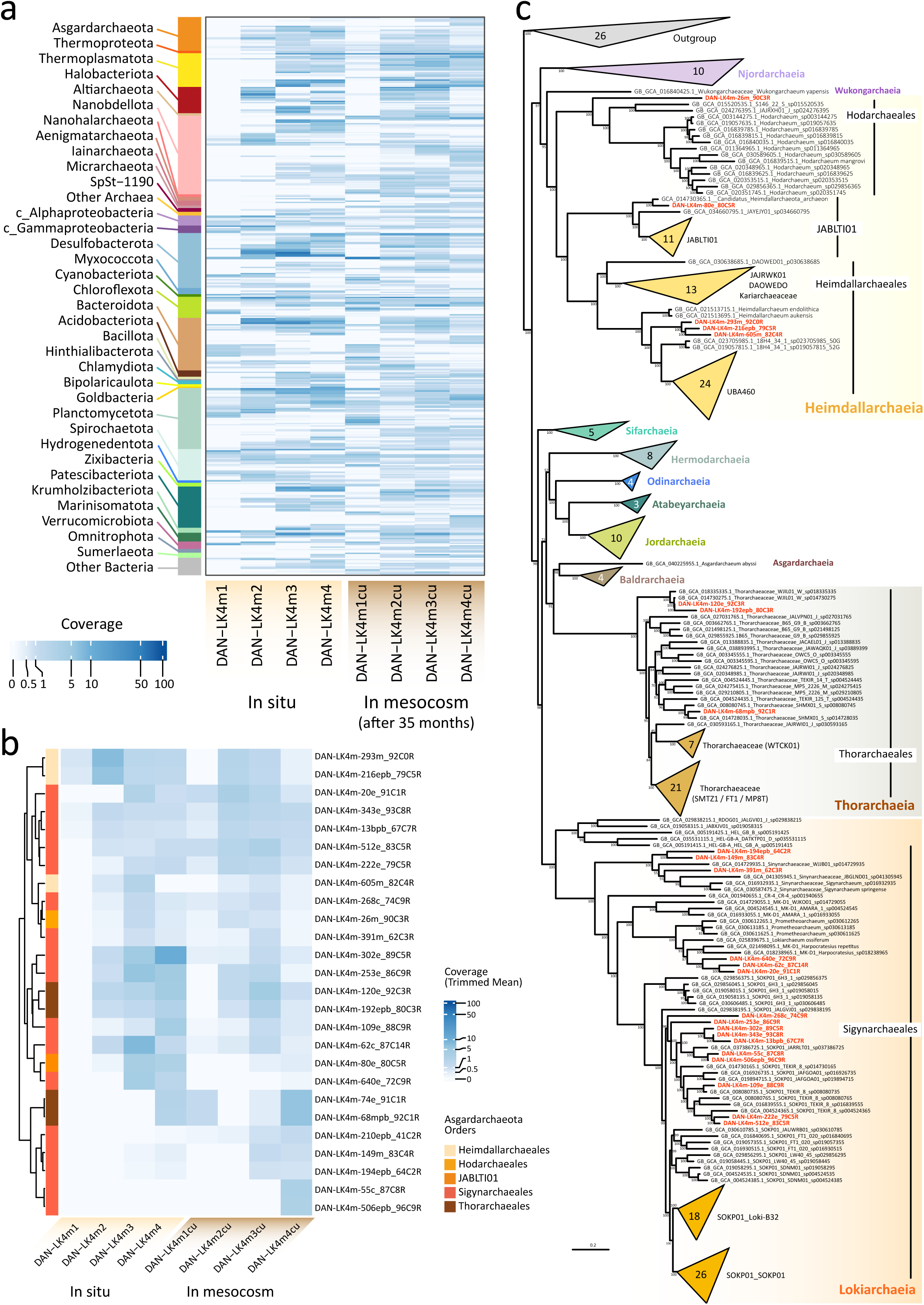
Distribution, relative abundance and phylogenetic affiliation of metagenome-assembled genomes (MAGs) in DAN-LK4 microbial mat metagenomes. **a**, Heatmap showing the distribution and coverage of all assembled MAGs in the different metagenomes grouped at the phylum level (detailed version in Supplementary Fig. 2). **b**, Detailed heatmap showing the distribution and relative abundance of asgardarchaeal MAGs. **c**, Maximum-likelihood multi-gene phylogenetic tree showing the position of the DAN-LK4 MAGs (red labels) within Asgardarchaeota. The tree was reconstructed with IQ-TREE (EAL+C60+G model; 297 taxa; 10,852 amino acid positions) and rooted with a selection of archaea belonging to the TACK supergroup. Scale indicates the expected number of substitutions per site. The full tree is shown in Supplementary Fig. 3. Detailed information about individual MAGs is given in Supplementary Table 6.

### Community structure across the redox gradient of natural and mesocosm-maintained microbial mats

After DNA purification from replicate mat layer subsamples, 16S rRNA gene fragments were amplified using prokaryote-specific primers^57^, massively sequenced and, after quality control, used to determine strictly defined amplicon sequence variants (ASVs). Of the 3,598 ASVs detected, 832 were archaeal and 60 were affiliated with Asgardarchaeota based on similarity comparisons with the SILVA^58^ database (Supplementary Tables 2-3). In parallel, we directly sequenced metagenomes for each DAN-LK4 microbial mat layer using Illumina (Supplementary Table 4). We then searched for a set of 15 Universal Single-Copy Genes (USCGs; essentially ribosomal proteins) that capture well the diversity and relative abundance of archaea, bacteria and eukaryotes^59^. We phylogenetically assigned USCGs to taxa by blasting them to reference genomes of the Genome Taxonomy Database (GTDB; release 226) and used normalized counts (Reads Per Kilobase per Million mapped reads; RPKM) of these markers to estimate relative abundances of taxa (Supplementary Table 5). To facilitate comparison between the metabarcoding (ASVs) and metagenomic (USCGs) datasets, we adapted the SILVA taxonomic assignment of 16S rRNA-based ASVs to the GTDB taxonomy (Supplementary Table 3).

The overall community structure inferred from 16S rRNA gene amplicon ASVs and metagenome-derived USCGs was diverse and overall congruent for both, in situ and mesocosm-maintained microbial mat layers (Fig.1c-d). The upper 1-cm layer of the microbial mat as sampled in situ (DAN-LK4m1) was dominated by Cyanobacteria, Chloroflexota, Pseudomonadota (Alpha- and Gammaproteobacteria), Bacteroidota, Planctomycetota and Spirochaetota, with Desulfurobacterota, Verrucomicrobiota and other bacterial and archaeal phyla in lower proportions. A substantial fraction of Chloroflexota (e.g. Chloroflexales), Alphaproteobacteria (e.g. Rhodospirillales, Rhodobacterales) and Gammaproteobacteria (e.g. Chromatiales, Ectothiorhodospirales) corresponded to lineages of anoxygenic photosynthetic bacteria, likely using the abundant H_2_S as electron donor for photosynthesis. Together with cyanobacteria, photosynthetic taxa represented about one third of the surface community members (Fig.1c-d; Supplementary Tables 3 and 5). Deeper, anoxic layers (DAN-LK4m2-m4) displayed more similar patterns between them, with Desulfurobacterota, Bacteroidota and Planctomycetota as dominant bacterial taxa, accompanied by Bacillota in bottom layers. Pseudomonadota were still present in DAN-LK4m2 but in much lower proportions than in the surface layer, and the presence of non-photosynthetic taxa in this clade increased, e.g. members of the sulfur-oxidizing chemolitoauthotrophic Halothiobacillales^60^ (Supplementary Tables 3 and 5). Remarkably, archaea, which were diverse, considerably increased their proportion in deeper layers, reaching up to one third of the total community. Four archaeal phyla dominated: Halobacteriota, including classical Halobacteria but also Archaeoglobales and Methanosarcinales, Thermoplasmatota (several classes, notably DHVEG-1), Nanobdellota and Asgardarchaeota. Halobacteriota were more abundant in DAN-LK4m2 but their proportion seemed to increase in the deepest sampled layer, partly due to the presence of methanogenic members. Halobacteriota comprised a wide variety of anaerobic family members including sulfur-respiring lithoheterotrophic archaea (e.g. *Halodesulfurarchaeum*)^61^. Thermoplasmatota were only present in the deepest layers (DAN-LK4m3-m4). Nanobdellota and Asgardarchaeota slightly increased with depth, the latter representing up to 5% of the total community (Fig.1c-d).

After almost three years of incubation under mesocosm conditions, the community composition exhibited changes but major dominant phyla remained relatively preserved in deep anoxic layers (Fig.1c-d, right panels). The most remarkable changes corresponded to the increase of Desulfobacterota alongside a strong decline of Pseudomonadota in the surface layer (DAN-LK4m1cu), as well as the colonization and spread of Thermoplasmatota in DAN-LK4m2cu and even, albeit in minor proportions, DAN-LK4m1cu. Collectively, archaea increased their proportions to more than 45% of the community in anoxic layers. Asgardarchaeota slightly increased their proportion in the upper cm but maintained similar abundance profile across depth, increasing from surface to deep layers, and reaching up to 5-6% of the total community members in the bottom layer. Although part of the microbial mat community seemed relatively maintained in the mesocosm over time, our results indicate that photosynthetic members were declining, as they represented around one fifth of the total community in the surface layer (Fig.1c-d; Supplementary Tables 3 and 5). While changes in the environmental conditions (different light quality, lack of continuous geothermal fluid input, slightly lower pH) might explain this evolution, it is also possible that the whole system is evolving towards more degradative, anaerobic heterotrophic processes; or both.

### Metabarcoding versus metagenome profiling of Asgardarchaeota

We investigated the finer composition of Asgardarchaeota in the DAN-LK4 microbial mats. The relative abundance of 16S rRNA-based ASVs and metagenome-derived USCGs belonging to this phylum was similar across depths and time. However, at finer taxonomic resolution, we observed differences i) between the two different marker sets and ii) over time (Fig.1e). Available “universal” primers for prokaryotic 16S rRNA genes are not inferred to amplify genes from the different asgardarchaeal classes equally well^29,62^. Consistent with this, the selected primer pair (U515F/926R), despite seemingly having broader coverage^29^, largely failed to amplify Thorarchaeia and, to a lesser extent, Heimdallarchaeia 16S rRNA genes, while successfully amplifying Lokiarchaeia genes (Fig.1e). In addition to this amplification bias, we encountered a problem affecting 16S rRNA gene classification in databases. To properly assign 16S rRNAs to the correct Asgard taxa, we reconstructed a comprehensive phylogenetic tree including reference sequences retrieved from GTDB MAGs and described asgardarchaeal species genomes, together with our amplicon sequences as well as 16S rRNA gene sequences identified in our MAGs (see below; Supplementary Fig.1). We identified several instances of 16S rRNA amplicon sequences misassigned based on SILVA SSU 138.2, e.g. Lokiarchaeia misassigned to Odinarchaeia, or Hodarchaeales to Lokiarchaeia. We also detected two cases of 16S rRNA gene-MAG misassignment: the *Sigynarchaeum springense*, retrieved from hot spring sediment^63^, and a related MAG (GB_GCA_014729935.1, classified as Sigynarchaeaceae) possessed 16S rRNA genes that formed a clade branching sister to the Hodarchaeales, far from sequences of the lokiarchaeal order Sigynarchaeales, to which the MAGs belong based on multiple protein-coding genes (GTDB; clade indicated with dashed lines in Supplementary Fig.1). Interestingly, we retrieved a MAG branching in a similar position sister to Hodarchaeales (DAN-LK4m-26m_90C3R, see below), which suggests that this clade of 16S rRNA sequences that also included DAN-LK4 ASVs (labeled “Heimdallarchaeia_unclassified” in Fig.1e), might correspond to the same clade and be bona fide, albeit divergent, Hodarchaeales. We used this 16S rRNA gene phylogenetic tree to correct the affiliation of our amplicon sequences (Supplementary Table 3, highlighted in red). Although the proportion of Heimdallarcheia detected by metabarcoding was lower than that inferred from USCG counts, we successfully amplified genes of the three known Heimdallarchaeia orders (Heimdallarchaeales, Hodarchaeales, JABLTI01).

Based on metagenomic inference, Heimdallarchaeia of the order Heimdallarchaeales were relatively abundant in upper layers of the in situ mat, clearly dominating DAN-LK4m2. A small fraction was detectable in the upper, oxygen-containing layer, confirming their oxygen tolerance^30,31,33^. Hodarchaeales and JABLTI01 were rarer taxa, but seemed to be relatively more abundant in DAN-LK4m3-m4. Lokiarchaeia clearly dominated these two deep layers, Thorarchaeia increasing in the deepest one (Fig.1e). After 35 months in the mesocosm, the proportion of Heimdallarchaeia decreased while Thorarchaeia became more abundant in all layers.

### Metagenome-assembled genomes across the redox zonation

To get insight into the functional potential and redox distribution of dominant microbial species in the DAN-LK4 microbial mats, we reconstructed MAGs. To facilitate assembly, we co-assembled sequences from the different layers of the in situ and mesocosm-maintained microbial mats, i.e. DAN-LK4m1-m4 and DAN-LK4m1cu-m4cu, respectively. To facilitate scaffolding, in addition to short reads, we used a composite dataset of long reads (PacBio) from anoxic mat layers (Supplementary Table 4). We assembled a total of 638 MAGs (Supplementary Table 6), from which we retained 445 high-quality, dereplicated ones (157 archaeal, 288 bacterial) for downstream analyses. While the vertical distribution of archaeal and bacterial taxa based on MAG relative abundance matched rather well that inferred from USCGs in terms of phyla, specific MAG spatial preferences were observed across layers (Fig.2a; Supplementary Fig.2). Archaeal MAG occurrence and relative abundance were lower in surface layers, increasing with depth, especially in the native DAN-LK4 mat. Patescibacteriota and many Desulfurobacterota and Planctomycetota MAGs exhibited similar trends. Reflecting the evolution of the mesocosm-maintained mat community, its finer MAG composition differed from that of the indigenous mat, notably in the two upper layers (DAN-LK4m1cu-m2cu). Some degree of homogenization of the microbial community, as seen through MAGs, across anoxic layers was also observed (Fig.2a). Likewise, asgardarchaeal MAGs exhibited spatial preferences across the redox gradient. These differences appeared slightly more marked in the indigenous Catherine volcano mat, where members of some Heimdallarchaeales were more abundant in intermediate layers close to the surface (DAN-LK4m2), whereas several Thorarchaeales and Sigynarchaeales (Lokiarchaeia) clearly preferred the deepest layers (Fig.2b).

We assembled MAGs for five orders of three Asgardarchaeota classes (Heimdallarchaeia, Lokiarchaeia and Thorarchaeia). Although Baldrarchaeia and Jordarchaeia were detected in DAN-LK4 metagenomes, they were in minor proportions, insufficient for MAG assembly. To ascertain the diversity of asgardarchaeal MAGs, we reconstructed a multi-gene phylogenetic tree using a maximum likelihood approach (Fig.2c; Supplementary Fig.3). We only included in this tree 24 high-quality non-redundant Asgardarchaeota MAGs that captured, based on average nucleotide identity (ANI), the specific diversity for this phylum (Supplementary Fig.4; Supplementary Table 7). Most MAGs branched within three families of the lokiarchaeal order Sigynarchaeales: Sigynarchaeaceae, MK-D1 (including *Prometheoarchaeum* and *Lokiarchaeum*) and SOKP01. Despite the higher relative abundance of Thorarchaeia in some mat layers, they were less diverse, belonging to the family Thorarchaeaceae (only three MAGs from two genera, WJIL01, SHMX01). Heimdallarchaeia, despite their lower abundance, covered a wider phylogenetic spectrum, with members branching within the orders Heimdallarchaeales, JABLTI01 and Hodarchaeales. Interestingly, the divergent MAG DAN-LK4m-26m_90C3R branched robustly as sister to previously known Hodarchaeales, defining at least a new family (Fig.2c). Unfortunately, although a correspondence is likely, a direct link with the 16S rRNA gene sequences mentioned above (Heimdallarchaeia_unclassified; Supplementary Fig.1) could not be established.

### Co-occurrence of Asgard archaea and other taxa

Asgardarchaeota showed a marked preference for the sulfate-reduction zone (SRZ), i.e. the area associated with H_2_S production along redox gradients (such as sediments or microbial mats, usually succeeded in deeper areas by a sulfate-methane transition zone, SMTZ). There, they broadly co-existed with co-dominant Desulfurobacterota, Bacteroidota, Planctomycetota and Thermoplasmatota, alongside less abundant taxa (Fig.1). Aiming to explore more specific shared preferences at lower taxonomic levels, we computed diverse co-occurrence networks based on normalized USCGs and 16S rRNA gene ASVs. Although the number of samples was low (n=8 metagenomes for USCGs; n=16 ASV datasets), these networks might reveal more specific neighborhoods and similar ecological preferences of Asgardarchaeota and other taxa. We first built a co-occurrence network based on USCGs using SparCC, given its better performance with low-sample compositional data^64,65^, and used it as primary reference for comparison. To ease visualization, we aggregated co-occurrences at the order level. The global co-occurrence network inferred in this way was complex (Supplementary Fig.5a), and we focused on the subnetwork containing asgardarchaeal nodes (Supplementary Fig.5b). Fig.3a summarizes positive correlations found between the different asgardarchaeal orders, proportionally distributed in mat layers, and different bacterial and archaeal taxa grouped at phylum level. Desulfurobacterota, Myxococcota, Bacillota and Thermoplasmatota exhibited the most connections with Asgardarchaeota, followed by Halobacteriota, Planctomycetota and others. Thorarchaeales, Heimdallarchaeales and Hodarchaeales correlated positively with more phyla than Sigynarchaeales, despite the latter being the most abundant and diverse Asgard order (Fig.2c). Fig.3a clearly shows the differential vertical distribution of Heimdallarchaeales, more represented in upper layers, as compared to Sigynarchaeales or Thorarchaeales. To visualize more specific co-occurrences, we extracted ego-networks showing asgardarchaeal orders and their positively and negatively correlated first neighbors at the order level (Fig.3b; Supplementary Fig.5c). Hodarchaeales displayed the highest number of positive correlations, mostly with deltaproteobacteria (Desulfurobacterota and Myxococcota) and Halobacteriota (both haloarchaea and methanogens), followed by Thorarchaeales. Not unsurprisingly, Desulfurobacterota members were found as positively correlated nodes in almost all ego-networks, and cyanobacteria, when present, always negatively correlated.

To refine our approach, we also generated co-occurrence networks based on both, USCGs and ASVs, using Spearman correlations with false discovery rate (FDR) correction, and a more exploratory SPIEC-EASI network. Correlation-based approaches capture both direct and indirect co-variation, whereas SPIEC-EASI aims to infer conditional relationships within a sparse graphical framework. Whereas SparCC produced the densest ego-networks using USCGs, the Spearman network retained fewer but stronger correlations (stricter correlation and significance thresholds were applied), and the SPIEC-EASI network was markedly sparser, consistent with its objective of inferring conditional dependencies and reducing indirect associations (Supplementary Fig.6). The Spearman and SPIEC-EASI networks obtained from ASV data were denser than USCG networks (Supplementary Figs.6-8). To explore correlation patterns at a finer level, we also plotted co-occurrences at MAG level (Supplementary Figs.9-10). Despite differences in network density, some consistent patterns emerged across methods. To integrate this information, we generated heatmaps showing the number of times than a given taxon pair was correlated positively or negatively across USCG networks (Fig.3c). In addition, consistency with the ASV networks was indicated on these heatmaps (asterisks). Given the amplification bias affecting 16S rRNA gene ASV data, the observation of a given pair also identified by USCG networks indicates a robust co-occurrence pattern; however, the absence of such observation does not indicate weaker potential interactions.

**Fig. 3.**
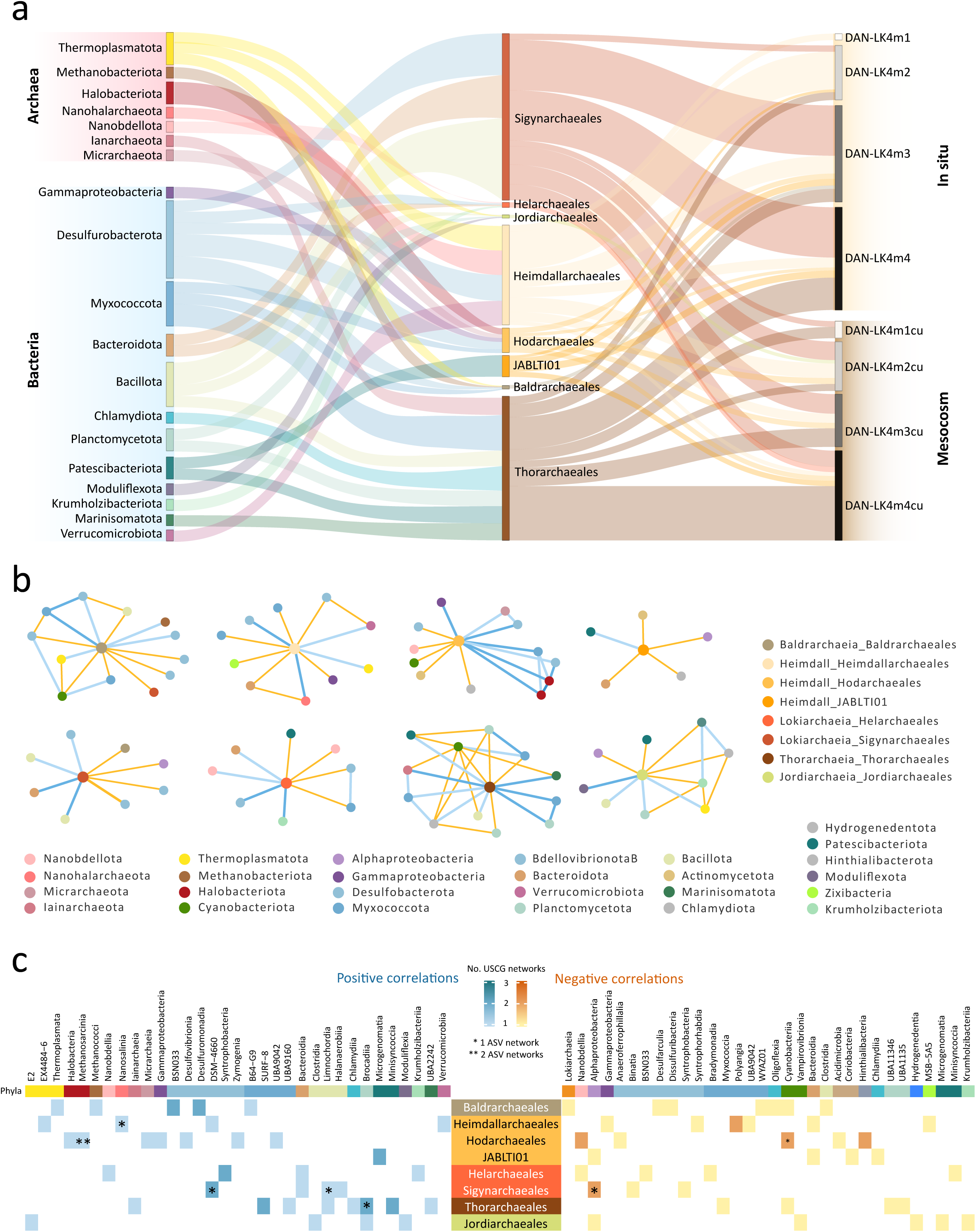
Correlations of Asgardarchaeota with other microbial lineages in DAN-LK4 microbial mats inferred by co-occurrence network analyses. **a**, Sankey plot showing positive correlations of asgardarchaeal orders based on normalized USCGs (RPKM) inferred by SparCC alongside their distribution across the redox gradient in the indigenous and mesocosm-maintained microbial mat. The size of boxes is proportional to the relative abundance of Asgardarchaeota (center) and the number of links with prokaryotic taxa (left) in co-occurrence networks. The relative abundance of the different asgardarchaeal orders is indicated by the thickness of links connecting with the different mat layers (right). **b**, Ego-networks of asgardarchaeal orders extracted from co-occurrence networks built with SparCC. Positive correlations are shown by blue edges, with darker shades corresponding to higher ⍴ values (e.g., ⍴ ≥ 0.7 was the darkest navy, whereas ⍴ near 0.3 was the lightest blue); negative correlations are shown as orange-yellow edges. Original and detailed networks are shown in Supplementary Fig.5. **c**, Heatmaps summarizing co-occurrence pairs observed across five different co-occurrence networks built from normalized USCGs (SparCC, Spearman correlations, SPIEC-EASI) and 16S rRNA gene-based ASVs (Spearman correlations, SPIEC-EASI). Blue, positive correlations; orange-yellow, negative correlations. The color shade indicates the number of times that a given pair is observed using USCGs (1–3); asterisks indicate additional recovery in one or both ASV networks. Co-occurrence networks were done at the order level; taxa co-occurring with asgardarchaeal groups were grouped at phylum level (except for Pseudomonadota, at class level; a-b) or class levels (c) to facilitate visualization. Ego-networks of Asgardarchaeota inferred using Spearman correlations and SPIEC-EASI based on USCGs and ASVs are shown in Supplementary Figs.6-7.

Asgardarchaeal orders, particularly Thorarchaeales, Sigynarchaeales and Baldrarchaeales, were most frequently observed to correlate positively with orders of the two deltaproteobacterial taxa Desulfurobacterota and Myxococcota (Fig.3c). Other pairs recurrently observed were Thorarchaeales and Brocadiia (Planctomycetota) or Minisyncoccia (Patescibacteriota). Intriguingly, the heimdallarchaeial order JABLTI01 only displayed a positive correlation with Microgenomatia, also member of the Patescibacteriota. Members of this bacterial phylum have reduced genomes and are inferred to be obligate symbionts/parasites of other bacteria^66,67^. However, across-domain episymbioses with methanogenic archaea have been recently described^68,69^, opening the door to possible episymbioses with other, including Asgard, archaea. Less unexpectedly, a recurrent correlation of Heimdallarchaeales with Nanosalinia, a member of the episymbiotic Nanohalarchaeota (DPANN archaea) was observed. Although Hodarchaeales exhibited positive correlations with diverse taxa, they most recurrently co-occurred with methanogenic Halobacteriota (Methanosarcinales). Among negative correlations, the most frequently observed were Hodarchaeales with Cyanobacteriia and Sigynarchaeales with Alphaproteobacteria (Fig.3c), highlighting the preference of these archaeal taxa for deeper layers in the microbial mats.

The inferred network associations should be interpreted cautiously as hypotheses of closer co-occurrence or shared environmental preferences. Nevertheless, the convergence of specific associations across independent inference methods might highlight candidate taxa for potentially closer, symbiotic and/or otherwise, ecological interactions.

### Asgard archaea and metabolic patterns across the redox gradient

To investigate dominant energy and carbon metabolism pathways in the DAN-LK4 microbial mats and their variation across the vertical redox gradients, we searched for a large collection of diagnostic metabolic genes (Supplementary Table 10), calculated their relative abundance across metagenomes (RPKMs) and visualized them in a heatmap (Fig.4a; Supplementary Fig.11 for a more detailed distribution of the corresponding genes). As expected, genes involved in oxygenic and anoxygenic photosynthesis were more abundant in the upper (1 cm) layer. However, minor proportions of these genes were detected in deeper layers, consistent with some degree of preservation of the corresponding microbes (or their DNA) during their progressive burial in the mat. Accordingly, genes for the complete Calvin cycle were dominant in the surface layer of the indigenous and the mesocosm microbial mats. Dissimilatory sulfate-reduction was widespread, being dominant in anoxic depths of the indigenous mat and equally distributed across layers in the mesocosm-maintained mat, in agreement with the distribution of Desulfurobacterota, the dominant sulfate-reducers, in the same mat (Fig.1c-d). Methanogenesis, albeit present, was much less represented, being slightly more abundant in DAN-LK4m3 and DAN-LK4m3cu. Oxidative phosphorylation genes were slightly more abundant in surface layers, but were also well represented in anoxic layers, including terminal oxidases (*coxA, coxB, coxC*) and high-affinity oxidases (*cydA, cydB*), the latter being more abundant, consistent with aerobic respiration in hypoxic environments^70^. Even if, like in the case of photosynthesizers, some of these genes might be associated with burying, dormant microorganisms, the quantitative persistence of these genes in anoxic layers appears counterintuitive, and might open the intriguing hypothesis that some of these terminal oxidases could suboptimally use the unstable intermediate NO as potential electron acceptor in the absence of molecular oxygen. Nitric oxide is transiently formed during denitrification (NO₃⁻ → NO₂⁻ → NO → N₂O → N₂), which also seems an active process in these microbial mats (Fig.4a). Oxygen reductases are structurally and mechanistically similar to NO reductases^71^, and they likely evolved from NO-reducing ancestors^72,73^. Superoxide dismutases, involved in ROS detoxification, were also widespread, highlighting their broad distribution in both, aerobic and anaerobic organisms^74,75^.

We searched for potential correlations between asgardarchaeal orders and specific metabolic pathways across the redox gradient (Fig.4b; Supplementary Fig.12). We observed a strong significant positive correlation of Hodarchaeales with methanogenesis, consistent with abovementioned observations showing the co-occurrence with methanogenic archaea within the Halobacteriota (Fig.3). Interestingly, members of the Hodarchaeales brought into culture establish syntrophic interactions with methanogens^32^, suggesting that they may be preferred hydrogen sinks for members of this asgardarchaeal order in natural ecosystems. Sigynarchaeales also positively correlated with methanogenesis, albeit less strongly (Fig.4b). Heimdallarchaeales strongly and positively correlated with oxygen-tolerant type 1 hydrogenases (indicated by their large catalytic subunit HyaB) and, to a lesser extent, sulfate reduction. Among others, these are essential respiratory hydrogenases in some Desulfobacterota, coupling hydrogen oxidation to reduction of Fe(III), fumarate or other electron acceptors^76^. Indeed, Heimdallarchaeales also significantly correlated with sulfate reduction. Mirroring the positively correlated metabolisms with different asgardarchaeal clades, Hodarchaeales very strongly anticorrelated with genes involved in oxygenic and anoxygenic photosynthesis and essential cyanobacterial hydrogenases (HoxH). Likewise, Thorarchaeales significantly anticorrelated with oxygenic photosynthesis and the Calvin-Benson C fixation pathway, and Sigynarchaeales, with both, oxygenic and anoxygenic photosynthesis, although less strongly. This emphasizes the preference of these archaeal clades for the anoxic layers and stricter anaerobic processes.

**Fig. 4.**
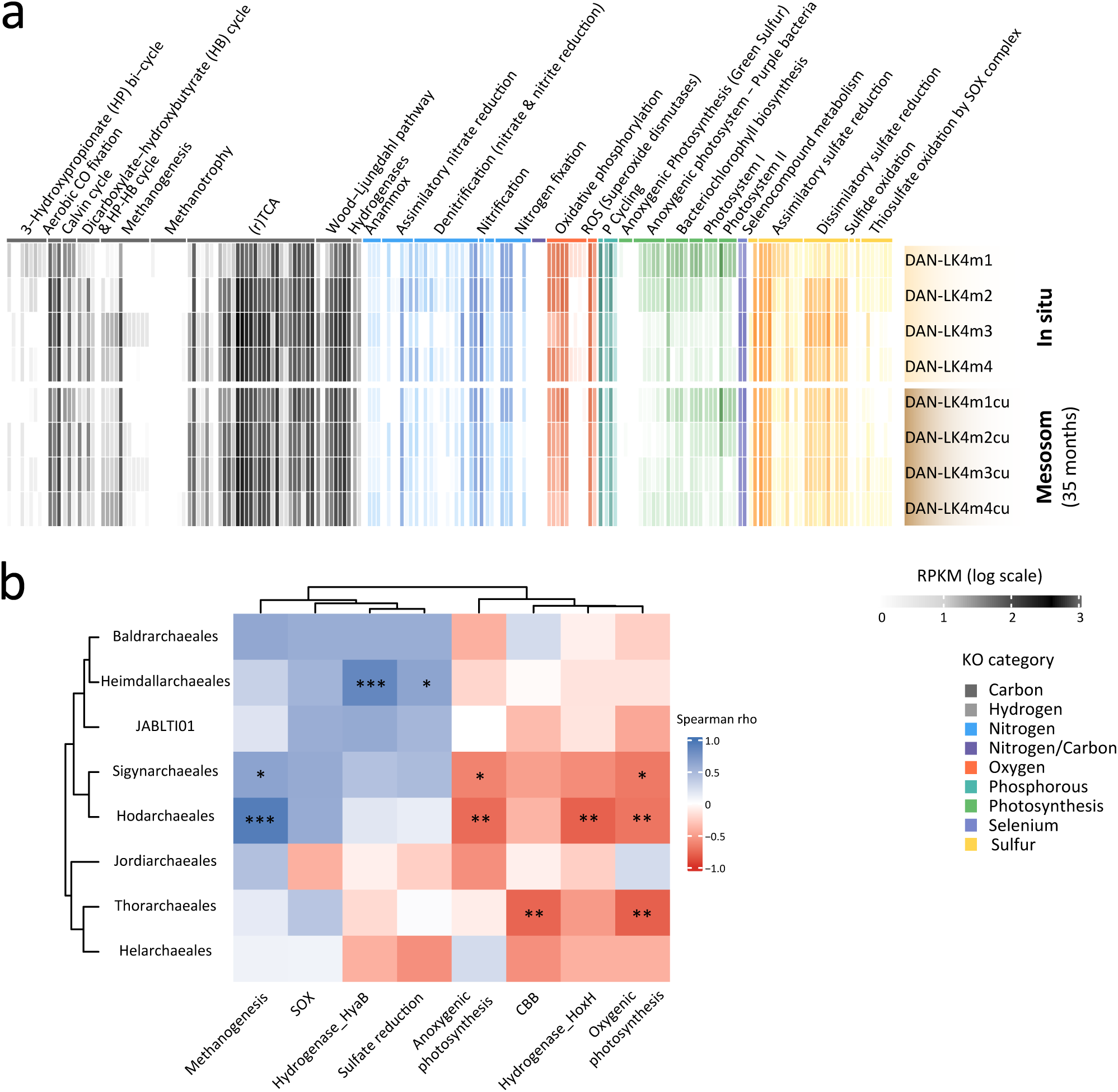
Redox distribution of metabolic pathways across DAN-LK4 microbial mat layers and correlations with asgardarchaeal taxa. **a**, Heatmap showing the relative abundance of genes involved in major metabolic pathways in metagenomes from different layers of the in-situ and mesocosm-maintained Catherine volcano lake microbial mats. A detailed version showing gene names is given in Supplementary Fig.11. **b**, Heatmap showing the sign and intensity of correlations between asgardarchaeal orders and different metabolic traits across the eight analyzed metagenomes (for a detailed version with ρ-values, Supplementary Fig.12a). SOX, sulfur oxidation; CBB Calvin-Benson-Bassham carbon fixation pathway. *** p<0.01, ** p<0.05, * p<0.10.

To explore potential metabolic complementarities and syntrophic interactions between Asgard archaea and other members of the community, we inferred the metabolic potential of representative MAGs for the Lokiarchaeia (Sigynarchaeales), Thorarchaeia (Thorarchaeales) and Heimdallarchaeia (Heimdallarchaeales and Hodarchaeales), screening the list of diagnostic genes alongside various transporter genes (Fig. 5; Supplementary Tables 10-12). We detected, in all cases, ABC transporters for oligopeptide uptake, a nearly complete Embden-Meyerhof-Parnas (EMP) glycolytic pathway, an incomplete tricarboxylic acid (TCA) cycle, a partial Wood-Ljungdahl (WL) pathway, diverse hydrogenases, and partial gene sets for the β-oxidation of fatty acids. The hodarchaeal MAG DAN-LK4m-26m_90C3R encoded multiple ABC transporters including several predicted to mediate the uptake of oligosaccharides such as maltose, cellobiose and chitobiose alongside oligopeptides, as did lokiarchaeial MAGs. Interestingly, DAN-LK4m-26m_90C3R and DAN-LK4m-74e_91C1R also possessed (nearly) complete pathways for nucleoside transport, nucleic acid degradation and salvage. Extracellular nucleosides might be converted into intracellular deoxypentose phosphates, thereby providing metabolic intermediates that may be indirectly linked to central carbon metabolism^77^. We also identified two types of NiFe hydrogenases in these two MAGs, a cytosolic Group 3b hydrogenase putatively involved in intracellular redox balancing via NAD(P)-dependent reactions, and a membrane-associated Group 4g hydrogenase likely linked to energy-conserving electron transfer (Fig.5a and c). While all our asgardarchaeal MAGs possessed genes involved in oxygen detoxification, sometimes in multicopy, none of them possessed genes involved in aerobic or nitrate respiration (Supplementary Table 11), suggesting that these archaea are anaerobes able to cope with oxidative stress. Collectively, Asgardarchaeota in the DAN-LK4 exhibited metabolic versatility, with the potential to utilize a broad range of organic substrates, and engage in hydrogen-mediated syntrophic interactions.

**Fig. 5.**
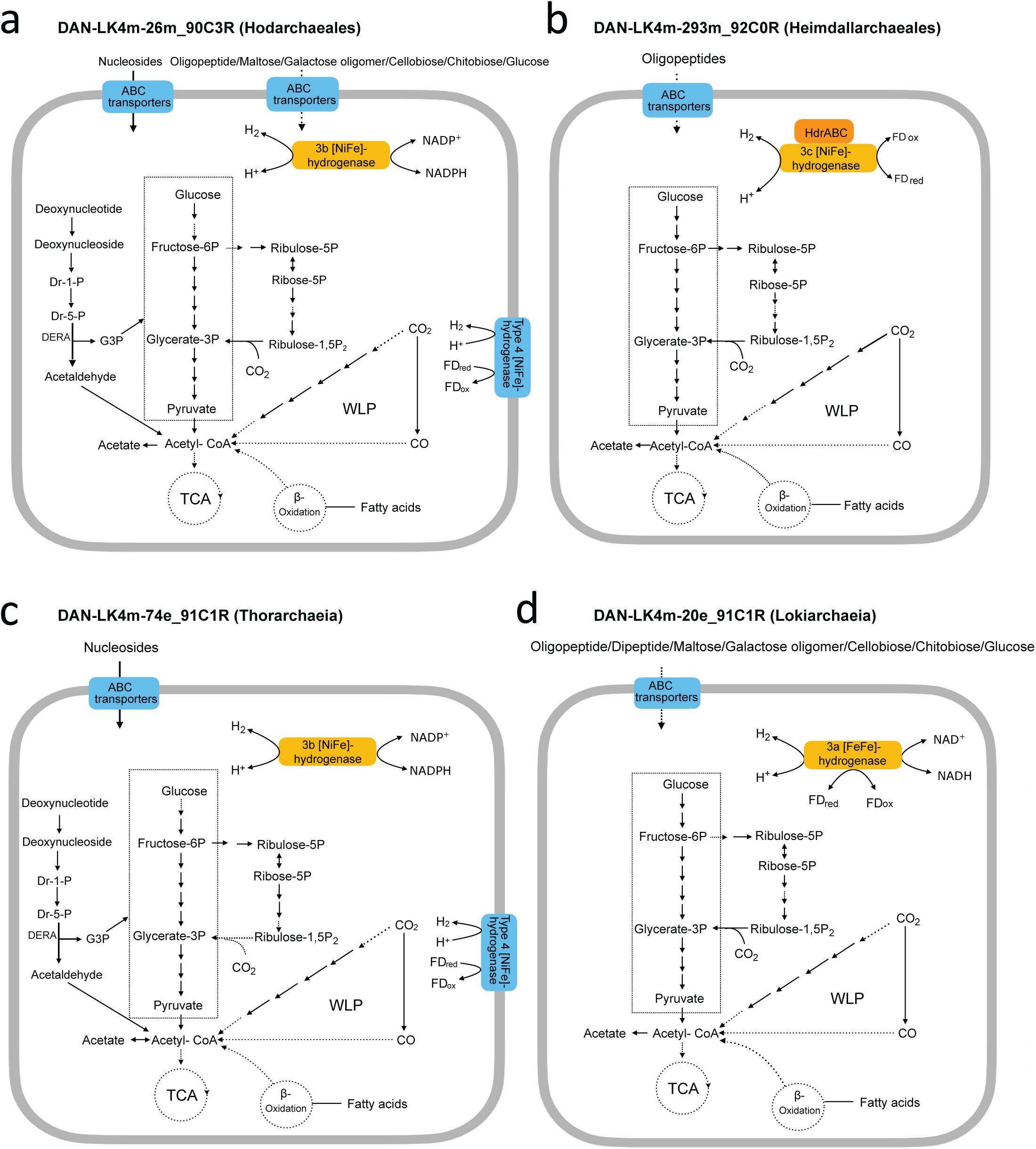
Major metabolic pathways of representative MAGs from DAN-LK4 microbial mats. **a**, DAN-LK4m-26m_90C3R belonging to the order Hodarchaeales (class Heimdallarchaeia). **b**, DAN-LK4m-293m_92C0R belonging to the genus UBA460, order Heimdallarchaeales (Heimdallarchaeia). **c**, DAN-LK4m-74e_91C1R belonging to the class Thorarchaeia. **d**, DAN-LK4m-20e_91C1R belonging to the class Lokiarchaeia. The arrows with solid black lines indicate genes present in the corresponding metabolic pathways, while those with dashed lines indicate absent genes, implying incomplete metabolic pathways. Dr1P, deoxyribose-1-phosphate; Dr5P, 2-deoxyribose-5-phosphate; DERA, deoxyribose 5-phosphate aldolase; G3P, glyceraldehyde-3-phosphate.

## Discussion

Eukaryotes likely evolved from the physical and genomic integration of an inter-domain symbiosis involving one Asgard archaeon and one or more bacterial partners^4,5^. This process occurred in early Proterozoic environments when atmospheric oxygen levels were much lower than today^45,46^. Precambrian redox-transition environments, such as microbial mats or shallow sediments, are favored cradles for eukaryogenesis^21^. There, the inherently benthic and ancestrally anaerobic Asgardarchaeota could stably interact with the facultatively aerobic alphaproteobacterial ancestor of mitochondria. Given that core metabolic functions are far more conserved than the organisms that carry them^78,79^ and that microbial communities are strongly subject to environmental selection^80,81^, similar core metabolic functions and interactions likely persist today in ecosystems resembling those where eukaryotes evolved. To apply this form of actualism^38^, we studied microbial mats from the saline, sulfidic and oxygen-poor lake (DAN-LK4) of the Catherine volcano, which bears conditions reminiscent of the late Archaean-early Proterozoic aquatic systems^48–50^. We combined 16S rRNA gene metabarcoding, metagenomic and genome-resolved analyses to characterize the community structure and functional potential across redox-gradient layers in the original microbial mat as well as a microbial mat maintained for almost three years in a laboratory mesocosm. We observed that, at phylum level, the overall community structure, including diverse Asgardarchaeota (up to ∼5% of the total community), was remarkably conserved between the two systems, especially in anoxic layers (Fig.1). The possibility of maintaining Asgardarchaeota over time in mesocosm opens interesting perspectives to study complex interactions in the laboratory, in ways complementary to more reductive studies based on antibiotic-treated enrichment cultures.

Distribution patterns of 16S rRNA gene-based ASVs, metagenome-derived USCGs, and MAGs in the two mats converged, despite temporal variation in lower-ranking taxa and experimentally confirmed primer bias against Thorarchaeia and, to a lesser extent, Heimdallarchaeia. Globally, Heimdallarchaeia, were more abundant in upper, more oxygenated layers, whereas Lokiarchaeia and Thorarchaeia preferentially thrived in deeper, anoxic layers. This confirms and extends previous observations showing that, compared to other Asgardarchaeota, Heimdallarcheia occupy more oxidized niches^30,31,35,82^, being aerotolerant^31,32^. However, within Heimdallarchaeia, only members of the Heimdallarchaeales exhibited a clear dominance in upper layers, significantly correlating with oxygen-tolerant hydrogenase genes (Fig.4), whereas members of the Hodarchaeales seemed to prefer more anoxic layers. Indeed, Hodarchaeales significantly correlated with methanogenic archaea and methanogenesis-related genes, typically occurring in the most anoxic zones, and anti-correlated with cyanobacteria and both, oxygenic and anoxygenic photosynthesis (Figs.3-4). Although we detected members of the Jordiarchaeia and Baldrarchaeia, they were relatively less abundant and we could not generate MAGs for them. Metabolic reconstruction suggests that DAN-LK4 Asgardarchaeota are versatile and can import and ferment oligopeptides and, frequently, diverse sugars. A divergent Hodarchaeales relative can also import and metabolize nucleosides. Several MAGs contain hydrogen-evolving hydrogenases (Fig.5). These metabolic features place Asgardarchaeota as intermediate in anaerobic, hydrogen-dependent processes, which are largely dependent on syntrophic interactions.

Sulfate-reducing bacteria, in particular Desulfurobacterota and Myxococcota^83^, are the most likely syntrophic partners for most Asgardarchaeota in these systems. Asgardarchaeota are well-distributed in the sulfate reduction zone (Fig.1), where these bacterial phyla are highly abundant. Co-occurrence networks also showed the highest number of significant correlations with deltaproteobacterial taxa (Fig.3). Sulfate-reducers and Asgardarchaeota usually co-occur also in other systems^14,29^. Although this is not direct proof, it is consistent with the long-recognized propensity of deltaproteobacterial sulfate-reducers to establish syntrophies with a variety of hydrogen-producing organisms^84–87^, including anaerobic protists^88,89^. Highly versatile, they can also be involved in mixed processes, shifting from sulfate-reduction to nitrate respiration and even aerobic respiration in some cases, depending on fluctuation of local conditions^90–92^. Finally, most Asgard archaea in culture are obligate syntrophs of Desulfobacterota species^19,42–44^. Furthermore, in cultures established from coastal microbial mats collected from the stromatolite-bearing Shark Bay (Australia), nanotubes connecting sulfate-reducing bacteria and asgardarchaeal cells indicate tight, indigenous symbiotic interactions^44^. It has been recently suggested that eukaryotes derived from an aerobic heimdallarchaeon that incorporated an aerobic alphaproteobacterium as endosymbiont, based on the detection of terminal oxidases and ROS-detoxifying enzymes in several heimdallarchaeal MAGs retrieved from oxygenated shallow sediments^31^. However, while we detected diverse genes involved in ROS detoxification in the DAN-LK4 asgardarchaeal MAGs, they consistently lacked any terminal oxidases. Given that this ecosystem is a better analog of early Proterozoic environments, and that eukaryogenesis started roughly 2 billion years ago, much before the atmosphere acquired its current oxygen level^45,46^, one might wonder instead whether the acquisition of those terminal oxidases by horizontal gene transfer in some modern Heimdallarchaeia did not postdate eukaryogenesis. Enzymes coping with ROS are broadly distributed across life domains and thought to have evolved early. It has even been suggested that reactive sulfur species (RSS) triggered the evolution of ROS antioxidants prior to the increase of oxygen levels^93^. A better knowledge of Asgard archaea ecology in more, and more diverse, Precambrian-like ecosystems should shed more light on the type and nature of symbiotic interactions that they entertained with bacteria at the dawn of eukaryogenesis.

## Methods

### Sampling and environmental parameters

Microbial mat samples were collected in January 2019 from the west shore of the Catherine volcano crater lake (numbered DAN-LK4 in a series of unnamed lakes sampled during the same expedition), north to the Erta Ale volcanic chain and south to the Danakil Depression, Afar region, Ethiopia (Supplementary Table 1; Fig. 1). The lake, located on volcanic malpais of extremely difficult access, was reached with the help of a helicopter (Tropic Air Kenya). Different physicochemical parameters (pH, temperature, conductivity, dissolved oxygen, redox potential) were measured in situ with a YSI Professional Series Plus multiparameter probe. Hydrochemistry data were generated from lake water filtered through 0.2 µm pore-diameter filters and collected in rubber-sealed glass bottles filled to the top to prevent prolonged contact with a fully oxygenated atmosphere. Major and trace elements were analyzed by Total Reflection X-Ray Fluorescence (TRXF; TXRF-8030c FEI spectrometer) and Inductively Coupled Plasma Mass Spectrometry (ICP-MS; Perkin–Elmer NexION 300XX instrument) at the SIDI Service (Servicio Interdepartamental de Investigación, Universidad Autónoma de Madrid). The content of major anions was analyzed by ionic chromatography (Flex 930 – Metrohm apparatus) at Qarbone 6napse (https://qarbone.com/). A microbial mat fragment of approximately 7 cm height, 18 cm width and 25 cm long was cut and collected in situ (Fig. 1a). A ∼1 cm-long section at one end of the mat was sliced in four different layers along depth as indicated in Fig.1a (DAN-LK4m1, ∼0-1 cm; DAN-LK4m2, ∼1-3 cm; DAN-LK4m3, ∼3-5 cm; DAN-LK4m, ∼5-7 cm) and fixed in 50-ml Falcon tubes with absolute ethanol. Fixed samples were subsequently stored at -20°C until further processing. The remaining mat fragment was immediately sealed in a plastic bag after removing most of the air from it. Back in the laboratory, this mat fragment was used to set up a small mesocosm in a transparent box placed in a culture room at ∼25°C under white light illumination (LED, 5,000K), with a daylight period of 12 h. Water from the lake (prefiltered by 0.2 µm pore-diameter filters) was used to cover the microbial mat to mimic as much as possible the local conditions. After 35 months of mesocosm incubation, we subsampled the microbial mat by taking a small core of ∼1 cm of diameter in the center of the mat throughout its entire depth. The core was divided in 4 layers (DAN-LK4m1cu to DAN-LK4m4cu) proportional to the original ones. This material was used to purify DNA for subsequent analyses. The oxygen (% saturation), pH and H_2_S (micromolar) vertical profiles of the long-term mesocosm were obtained using a Field MicroProfiling FMM system (UNISENSE, Denmark) equipped with specific 50-µm electrodes, calibrated according to the manufacturer’s specifications. Measurements were taken in triplicate (three different electrodes in three different locations in the mat) every 0.5 mm to a total depth of 45 mm. Deeper measurements could not be done due to the more conspicuous presence of mineral particles.

### DNA purification, 16S rRNA gene metabarcoding and metagenome sequencing

DNA was purified from biomass distributed in replicate tubes using the DNeasy Power Soil DNA Isolation Kit (Qiagen). In the case of fixed samples, the ethanol was removed and the samples rehydrated overnight at 4°C in the initial resuspension buffer of the kit. Final eluted DNA was resuspended in 10 mM Tris–HCl, pH 8.0 and stored at −20 °C. 16S rRNA gene amplicons (∼420-440 bp) were obtained from replicates (a, b) using the specific prokaryotic primers U515F (5’- GTGCCAGCMGCCGCGGTAA) and 926R (5’-CCGYCAATTYMTTTRAGTTT)^57^ tagged with different barcodes. Amplicons were multiplexed and then sequenced using 300-bp paired-end (PE300) MiSeq Illumina technology (Eurofins Genomics NGS Lab, Constance, Germany). For metagenomes, DNA from replicate extractions was mixed to reach a safe amount of DNA for sequencing. Sequences from the eight metagenomes (DAN-LK4m1 to DAN-LK4m4; and DAN-LK4m1_cu to DAN-LK4m4_cu) were obtained by NovaSeq 6000 S4 PE150 XP (Eurofins Genomics). To facilitate subsequent MAG assembly, the remaining DNA from layers below DAN-LK4m1/m1cu was mixed to construct a 10kb SMRTbell Express library and generate a metagenome encompassing several layers (DAN-LK4m_PacBio_M2M3M4M4cu) for long-read PacBiosequencing (Macrogen Europe).

### Sequence analysis and taxonomic profiling

16S rRNA gene amplicon sequences were analyzed using QIIME2 v.2021.4^94^. The “cutadapt demux-paired” function was used to demultiplex the different samples. We determined amplicon sequence variants (ASVs) using the DADA2 pipeline implemented in QIIME2 after denoising, dereplication, and chimera-filtering^95^. The optimal truncation parameters were first selected using FIGARO^96^ and then modified manually for optimal results (truncation of forward and reverse reads at 273 and 225 nt, respectively). Forward and reverse reads were merged to obtain complete denoised ASV sequences. Denoised sequences with one or more mismatches in the overlap region were discarded. ASVs were assigned to archaeal and bacterial taxa based on the SILVA^58^ SSU 138.2 release taxonomy using vsearch^97^ implemented in the QIIME2 pipeline^98^. The names of the different taxa were then converted to fit those of the GTDB taxonomy (https://gtdb.ecogenomic.org/) release 226 using an in-house script. Since, based on phylogenetic analysis, the SILVA assignment of Asgardarchaeota sequences was not reliable, we classified asgardarchaeal ASVs based on phylogenetic analysis (Supplementary Fig.1; see below). After sequence quality-filtering and reorientation, a total of 3,598 prokaryotic ASVs were obtained, 832 of which were archaeal ASVs (Supplementary Table 2). Sequences affiliating to chloroplasts, mitochondria or (rarely) eukaryotes were removed prior to the analysis. The relative abundance and taxonomic affiliation of ASVs are provided in Supplementary Table 3.

For metagenome sequence analysis, the quality of Illumina raw reads was initially inspected with FastQC^99^ v0.11.8. Subsequently, they were trimmed and filtered with Trimmomatic^100^ v0.39, using the following parameters adjusted based on the FastQC quality profiles: LEADING:3, TRAILING:3, MAXINFO:40:0.8, and MINLEN:36. Assembly was performed independently for the eight metagenomes using metaSPAdes^101^ v3.15.3, applying default options and a multi-k-mer strategy (k = 21, 25, 31, 35, 41, 45, 51, 55). Genes were predicted and annotated using Prokka^102^ v1.14.5 in metagenome mode, retaining only contigs ≥200 bp long. The PacBio long reads from the pooled M2-M4 layer metagenomes, were assembled using Flye^103^ v2.9.1 with a k-mer size parameter of 15, retaining only contigs ≥10 kbp in length. This dataset was used to generate longer contigs for improved genome reconstruction. Metagenome statistics are provided in Supplementary Table 4. To estimate the relative abundance of microbial taxa within the eight Illumina-sequenced metagenomes, we used a core set of 15 Universal Single-Copy Genes (USCGs)^59^ defined by their Pfam domains as annotated with the Pfam-A database v3.1b2 using HMMER hmmsearch^104^ v3.2.1. The resulting protein sequences were compared against representative genomes from GTDB r226 and NCBI RefSeq r232 via BLASTp v2.9.0. For each marker gene, taxonomy was assigned based on the top hit meeting minimum thresholds of ≥35% identity and ≥70% alignment coverage. To calculate the relative abundance of the different taxa, USCG nucleotide sequences were indexed with Bowtie2^105^ v2.3.5.1, and raw reads were aligned to them. Read mappings were filtered using Samtools^106^ v1.9, and gene-level abundances were calculated as Reads Per Kilobase per Million mapped reads (RPKM) via custom Perl scripts. RPKM values for each gene were then averaged across all USCGs assigned to the same taxon to obtain normalized abundance estimates at different taxonomic levels (e.g., phylum, class, order; Supplementary Table 5). Community composition plots were generated using *ad hoc* R^107^ scripts with the ggplot2 package^108^. Taxa comprising <0.5% of total abundance were grouped under “Other Bacteria” or “Other Archaea.”

### Reconstruction of metagenome-assembled genomes (MAGs)

The eight Illumina metagenomes were co-assembled using the same metaSPAdes parameters described above for binning purposes. The resulting dataset, referred to as DAN-LK4m, was subjected to genome binning with the anvi’o platform^109^ (developer version). Within anvi’o, we integrated outputs from four automated binning tools: MetaBAT2^110^, MaxBin2 ^111^, Binsanity^112^, and CONCOCT^113^. Bins were dereplicated using DAS Tool^114^ v1.1.6, and manually refined using anvi-refine. MAG quality (completeness and redundancy) was evaluated within the anvi’o environment. In parallel, we used PacBio contigs to help MAG assembly and binning. PacBio contigs larger than 1,000 bp were collectively treated as a co-assembly and binned with the anvi’o pipeline with the same parameters as above, mapping the 8 sets of Illumina reads back to it to estimate differential coverage. After dereplicating the Illumina and PacBio (co)assemblies, we obtained a total of 445 high-quality MAGs (157 archaeal, 288 bacterial) and excluded 193 MAGs displaying poorer metrics from the rest of the analyses (Supplementary Table 6). The taxonomic affiliation was carried out with GTDB-Tk^115^ v2.4.1 on GTDB release r226. 16S rRNA genes were identified with ssu-align^116^ v0.1.1, and predicted genetic codes were inferred using Codetta2^117^ v2.0 (Supplementary Table 6). Genes in the high-quality MAGs were annotated with Prokka^102^ using tailored parameters optimized for the codon usage and taxonomy of each MAG. MAG coverage across metagenomes was calculated by mapping each set of metagenomic reads to individual MAGs with CoverM^118^ v0.7.0. The distribution of the MAGs across the 8 different metagenomes was visualized with an in-house R script.

### Phylogenetic analyses

To confidently classify asgardarchaeal 16S rRNA gene sequences, we aligned our amplicon sequences with MAG-associated full-length 16S rRNA gene sequences from this study (6 MAGs) and from a wide representation of reference GTDB MAGs, including described Asgardarchaeota species, (393 total taxa). Nucleotide sequences were aligned using MAFFT^119^ with the accurate L-INS-i algorithm; the --adjustdirection option was used to automatically detect and correct sequence orientation prior to alignment. The untrimmed alignment (2,495 nucleotide positions) was used to reconstruct a phylogenetic tree using a maximum likelihood approach implemented in IQ-TREE^120^ v3.0.1 with the GTR+G4 substitution model. Node support was estimated using 1,000 SH-aLRT replicates and 1,000 ultrafast bootstrap replicates. The resulting phylogeny was used to refine the taxonomic annotation of ASVs inferred from SILVA. When an ASV was consistently placed within a well-supported clade composed of reference sequences assigned to a different taxon, the ASV taxonomy was revised to match the phylogenetic placement supported by the surrounding reference sequences. To place asgardarchaeal MAGs in a phylogenetic tree, we retrieved the sequences of 53 conserved proteins^16^ from our MAGs and representative MAGs across archaea (297 taxa). Each dataset was aligned using MAFFT-L-INS-i; ambiguously aligned positions were trimmed using TrimAl^121^. Trimmed alignments were concatenated into a supermatrix (297 taxa; 10,852 amino acid positions) and the resulting multi-gene alignment was used for phylogenetic reconstruction with IQ-TREE v3 under the EAL+C60+G model; 1,000 ultrafast bootstrap replicates were used to assess branch statistical support.

### Co-occurrence networks

To infer microbial co-occurrence networks from USCGs, we applied three approaches using as input their normalized abundance (RPKMs) aggregated at the order taxonomic level. Rare taxa (< 1% per metagenome) were excluded, except for Asgardarchaeota. First, we used SparCC (Sparse Correlation for Compositional data)^64^ to calculate correlation coefficients (⍴) and associated p-values. Ten iterations were used to estimate the median correlation of each pair, and their statistical significance was determined using 500 bootstrap iterations. To minimize the inclusion of spurious correlations, raw p-values for all pairwise interactions were adjusted using the Benjamini–Hochberg false discovery rate (FDR)^122^ in an *ad hoc* R^107^ script. Only correlations with an adjusted p-value (or Q-value) ≤ 0.05 and |ρ| ≥ 0.3 were retained. Networks were built using *ad hoc* R scripts and visualized with the aid of the igraph package (http://igraph.org/) v2.1.4 and Cytoscape7 (https://cytoscape.org/) v3.10.3.

Second, we estimated order-level pairwise associations using Spearman rank correlations after a centered log-ratio (CLR) transformation (after adding a small pseudocount to handle zeros) to account for the compositional nature of the data. The statistical significance of correlations was assessed using permutation tests (5,000 per pair of orders) to generate a null distribution of correlation coefficients. Empirical p-values (proportion of permuted correlations with absolute values ≥ observed correlations) were adjusted for multiple testing using the Benjamini–Hochberg FDR procedure. Edges with correlation strength |ρ| ≥ 0.3, without FDR filtering, were retained for cross-comparison; for visualization in Cytoscape, we also retained edges with FDR ≤ 0.05, to highlight the strongest statistically supported associations (Supplementary Fig.6a). A third, exploratory, network was inferred using SPIEC-EASI (Sparse InversE Covariance Estimation for Ecological Association Inference)^123^ with the Meinshausen–Bühlmann^124^ neighborhood selection method (λ grid: nλ = 30, λmin ratio = 10⁻²; model selection was done by stability selection using the StARS criterion with 50 resampling replicates). This approach estimates sparse conditional dependency structures and reduces the influence of indirect correlations. Given the limited sample size, SPIEC-EASI results were interpreted cautiously and used primarily for comparison with correlation-based networks. The MB model was fitted using a range of regularization parameters, with model selection performed via stability selection. The resulting binary adjacency matrix indicates putative direct associations between orders. Ego-networks of Asgard orders were also extracted from SPIEC-EASI networks for visualization in Cytoscape (Supplementary Fig.6b). To aid biological interpretation, Spearman correlation coefficients computed on the same filtered dataset were mapped onto the edges of the SPIEC-EASI network, providing both the direction and magnitude of co-variation for each inferred association.

To construct co-occurrence networks of 16S rRNA gene ASVs, we grouped them at order level, removing rare orders (<1 % relative abundance per sample). The filtered matrix was then normalized using a Hellinger transformation to reduce the influence of dominant taxa and account for the compositional nature of the data. A co-variation network was constructed from these transformed data using Spearman rank correlations between orders (Supplementary Fig.7) and an exploratory network was inferred using SPIEC-EASI (Supplementary Fig.8), as above. Edge tables containing source nodes, target nodes, Spearman correlation coefficients (ρ), and corresponding adjusted p-values for all networks and network cross-comparison are available in Supplementary Tables 8-9. Finally, to compare co-occurrence patterns involving Asgard archaea across USCG and ASV networks, edges from the five networks were integrated using the USCG SparCC network as the primary reference, given its better performance with low-sample compositional data^64,65^. All Asgard-target pairs detected in the SparCC network were retained, with Asgard taxa resolved at order level and partner taxa aggregated at class level. For each pair, recovery was additionally scored across the two remaining USCG networks (Spearman and SPIEC-EASI; edges with |ρ| ≤ 0.3 were excluded across all networks prior to aggregation; for Spearman networks, edges with q-value > 0.1 were additionally excluded) and across the two ASV-based networks (the ASV Spearman network was filtered at the same q-value threshold). Positive and negative associations were treated as biologically distinct, and visualized in independent heatmaps coded in R with the ComplexHeatmap library^125^, where cell color reflects the number of USCG networks recovering the pair (1–3) and asterisks indicate additional recovery in one or both ASV networks (Fig.3c).

To visualize co-occurrence patterns at MAG level, we extracted the first neighbors from the SparCC network for each asgardarchaeal order. The resulting order-level neighborhoods were used to identify the corresponding individual MAGs: for each network node (Asgard order or partner order), all dereplicated MAGs assigned to that taxonomic order were retrieved from the MAG taxonomy table. MAG abundances across the eight metagenomes were obtained as trimmed mean coverage values computed with CoverM, as described above. The coverage matrix was visualized as a heatmap using the ComplexHeatmap R package, where rows corresponded to individual MAGs, grouped by Asgard order co-occurrence neighborhood, and columns corresponded to metagenomes ordered by depth.

### Metabolic inference

The general metabolic potential of metagenomes was evaluated via the prediction of KOfam^126^ HMMs (v2025) using HMMER hmmsearch^104^ v3.2.1. Hits were filtered by e-value (≤ 1e-20) and selecting only one KO per gene (the match with the best e-value). We searched for 191 diagnostic metabolic genes (Supplementary Table 10), calculated their relative abundance (RPKMs; as described above for USCGs) and visualised them in a heatmap generated with ComplexHeatmap^127^ in R. Genes of asgardarchaeal MAGs were annotated using DRAM^128^ against KEGG^129^, PFAM^130^, dbCAN^131^ and MEROPS^132^ databases with default parameters (Supplementary Table 11). Annotated genes on oxygen-related metabolism as well as nitrate and aerobic respiration were selected and presented in Supplementary Table 11. The metabolic pathways for selected representative MAGs was inferred after annotation by BlastKOALA^129^ on the KEGG database followed by the identification of the metabolic pathways using KEGG Mapper^133^. The annotations of the MAGs are provided in Supplementary Table 11. Putative hydrogenases were identified using HydDB^134^ with an e-value cutoff of 1e-20, along with the CxxC motifs in the sequences (Supplementary Table 12). Spearman rank correlations were calculated for normalized abundances of Asgard orders and metabolic genes: Calvin-Benson-Bassham CBB (RbcL, RbcS, PRK; K0160, K01602, K00855), methanogenesis (McrA; K00399), hydrogenase HoxH (K00436), hydrogenase HyaB (K06281), anoxygenic photosynthesis (PufM, PufL; K08929, K08928), oxygenic photosynthesis (PsbA, PsbB; K02703, K02704), sulfate reduction (DsrA, DsrB; K11180, K11181), and sulfide/sulfur oxidation (SoxA, SoxB, SoxX, SoxY; K17222, K17224, K17223, K17226). Spearman correlations were computed for all order-metabolism combinations (n=64 tests). Analyses were performed in R using the ComplexHeatmap package.

## Data availability

Metagenome Illumina sequences and metabarcoding data have been deposited in GenBank (National Center for Biotechnology Information) Short Read Archive with BioProject number PRJNA1368766, BioSample SAMN53369320.

## Code availability

Custom code for these analyses (*perl* and R scripts) is available at Gitlab (https://gitlab.com/DeemTeam/dan-lk4m).

## Supplementary Material

Supplementary Tables are available at FigShare (https://doi.org/10.6084/m9.figshare.31956027)

## Acknowledgements

We thank Olivier Grunewald for the organization of the field trip, Luigi Cantamessa and his Tigrayan/Ethiopian team for the in situ logistics, Jacques Barthélémy, Abdul Ahmed Aliyu and the Afar authorities for local assistance and the Afar police for providing security. We thank Jodie Belilla and Ana I. López-Archilla for, respectively, help with sampling and hydrochemical analyses, and Timmy Flowers (Tropic Air Kenya) for efficiently piloting the helicopter. The field trip organization (O.G.) received support from the Mamont Foundation. This work was supported by the Gordon and Betty Moore Foundation (P.L.-G., https://doi.org/10.37807/GBMF9739), the Agence National de la Recherche (P.L.-G., ABiSYM ANR-23-CE02-0016-01; DM, DArchFolds, ANR-22-CE02-0012), and the ERC-AdG grants Plast-Evol (D.M., 787904) and SYMBEK (P.L.-G., 101141745). The equipment used to perform the microelectrode measurements was funded by Université Paris-Saclay (AAP 2023 “Innovation pédagogique” and Department of Biology).

## Author contributions

P.L.-G. and D.M. conceived the study, established mesocosms, organized the scientific field trip and collected samples and metadata. A.G.-P. analyzed metagenomic data with bioinformatic support by P.D. M.I. carried out metabarcoding analyses with the help of A.G.-P. and P.D. and measured redox profiles. Y.Z. assembled PacBio data. A.G.-P. and A. S. conducted network analyses. A.G.-P. and L.L. analyzed metabolic pathways in assembled metagenomes and MAGs. J.M.L.-G. and K.B. helped with sampling, georeferencing and hydrochemical interpretation. A.S., L.E and A.G.-P. carried out phylogenetic analyses. P.L.-G. wrote the manuscript. All authors read, commented and approved the manuscript.

## Competing interests

The authors declare no competing interests.

## Notes

### Competing Interest Statement

The authors have declared no competing interest.

### Summary of Updates

This version of the manuscript has been revised to slightly update the text, clarifying some aspects, and including minor changes in some figures.

