## Supplementary Figures for "Redox distribution of Asgard archaea and co-occurring taxa in microbial mats from an early Proterozoic ecosystem analog" (from another version of this preprint: version 1, 2026-03-21)

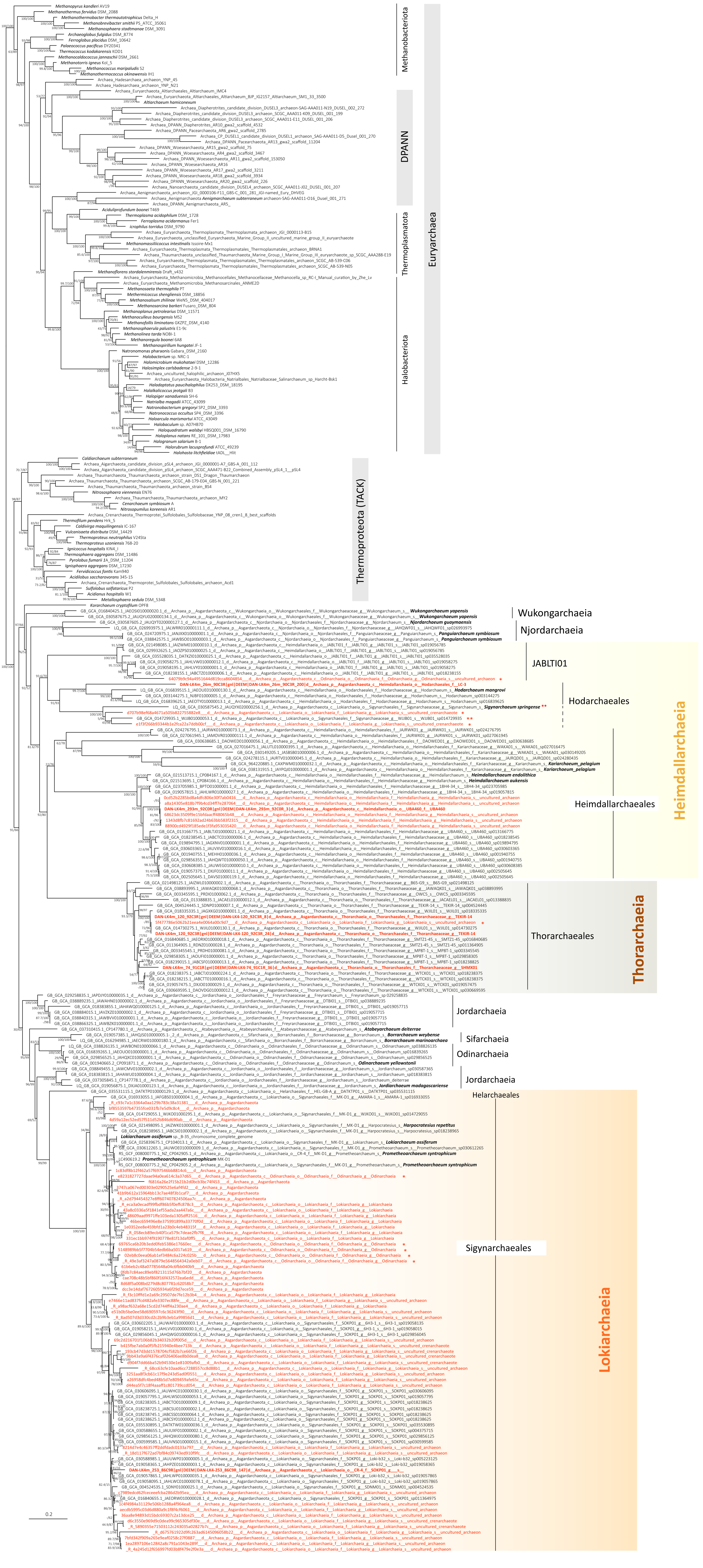


**Supplementary Fig.1. Phylogenetic tree showing the phylogenetic position of 16S rRNA gene sequences of Asgard archaea from the DAN-LK4 microbial mats.** Amplicon sequences from this work are shown in color. MAG-associated 16S rRNA genes from this work are shown in color and bold. Note that the phylogenetic assignation done with SILVA (*) or the affiliation of 16S rRNA genes associated with MAGs (**) does not always correspond to the correct Asgard archaeal group, indicating problems of sequence misassignment in (some) assemblies and sequence databases.

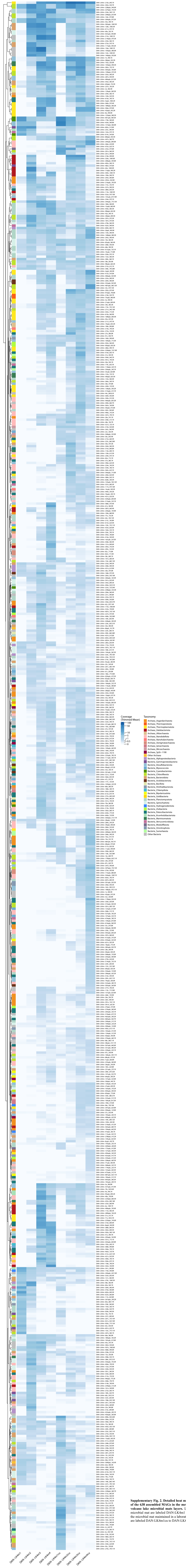


**Supplementary Fig. 2. Detailed heat map showing the distribution of the 638 assembled MAGs in the metagenomes of the Catherine volcano lake microbial mats layers.** Layers from the in-situ fixed microbial mat are labeled DAN-LK4m1 to DAN-LK4m4; those from the microbial mat maintained in a laboratory mesocosm for 35 months are labeled DAN-LK4m1cu to DAN-LK4m4cu.

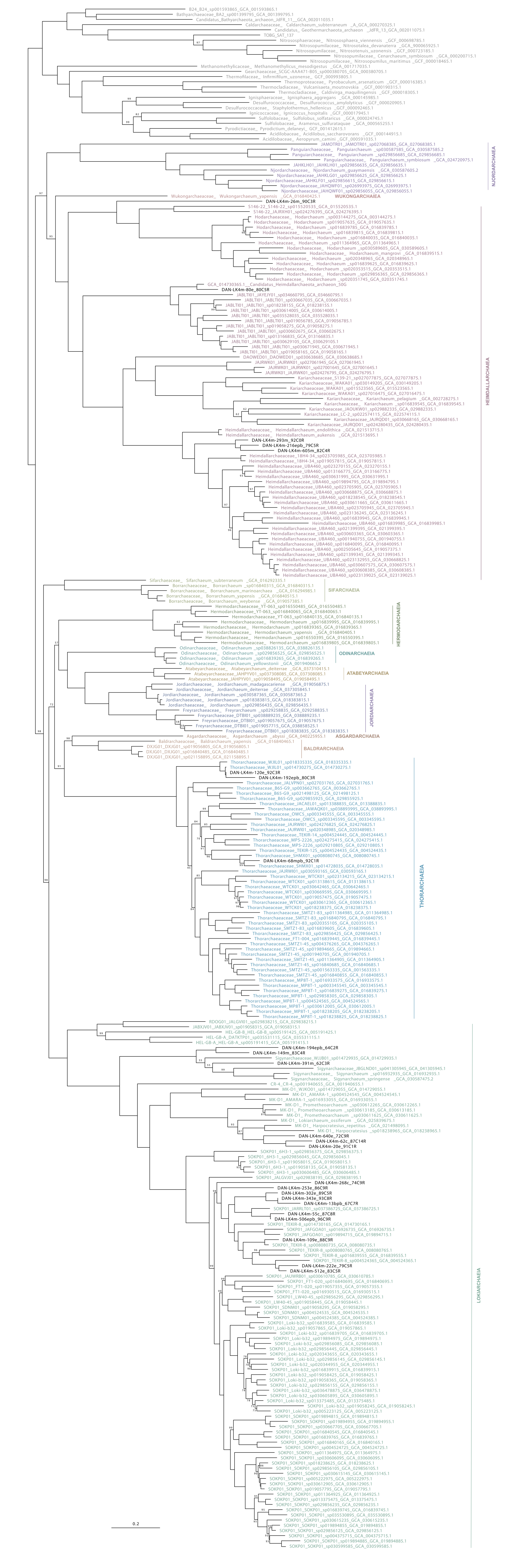


**Supplementary Fig.3. Rooted phylogeny of Asgard archaea highlighting the placement of Lake DAN-LK4 microbial mat MAGs.** The tree was estimated by Maximum Likelihood from a concatenated dataset of 53 proteins (297 taxa, 10852 positions), under the LG + EAL + G4 model. All branches have 100% ultrafast bootstrap support unless otherwise specified. Each leaf contains order, family, genus and species information according to the Genome Taxonomy DataBase, as well as the genome accession number. Classes are indicated near each color-coded clade. MAGs described as part of this work are displayed in bold. Scale indicates the expected number of substitutions per site.

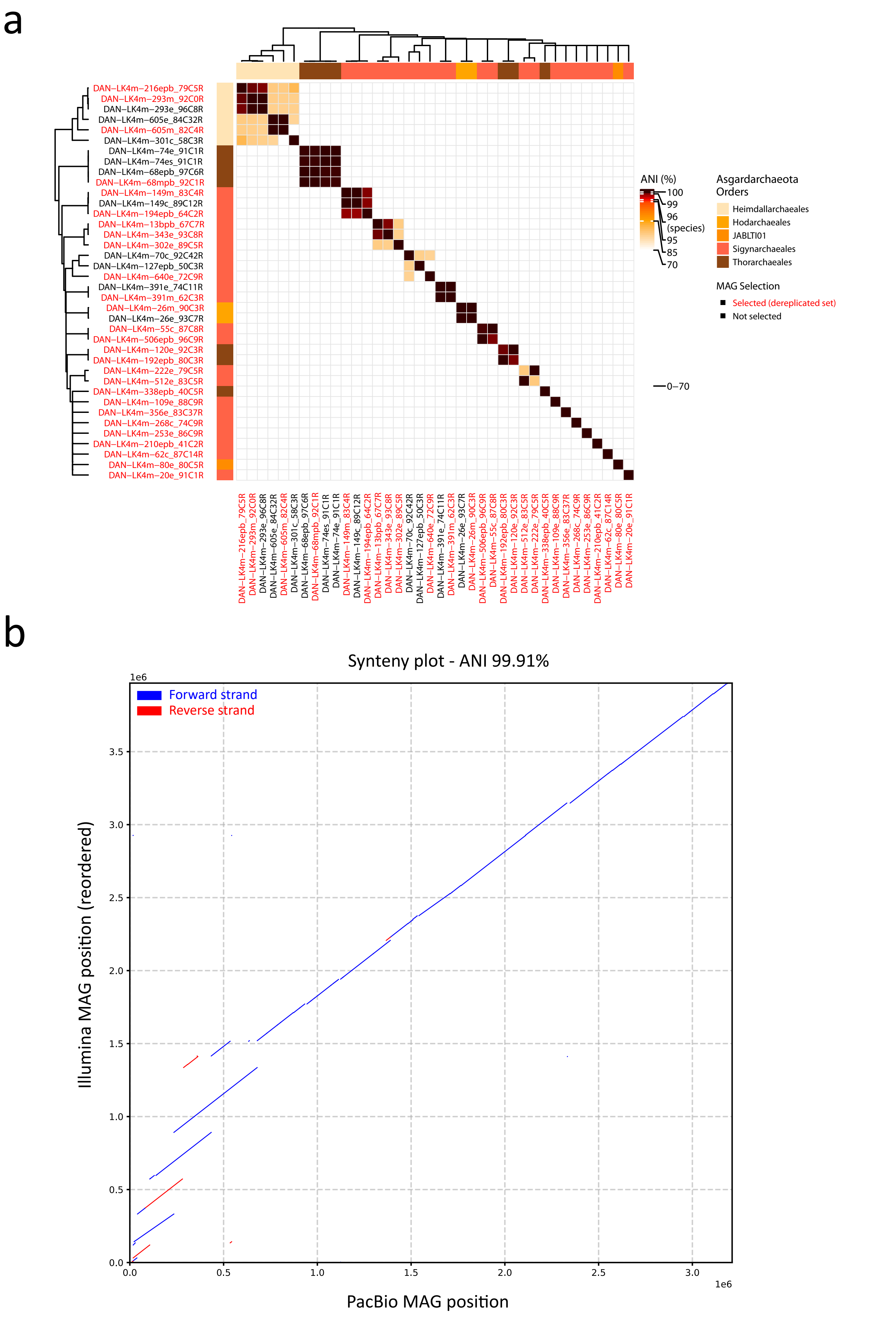


**Supplementary Fig. 4. Average nucleotide identity (ANI) of Asgard archaea MAGs.** a, Cluster analysis of asgardarchaeal MAGs according to their phylogenetic relatedness. b, Synteny plot of the Thorarchaeia MAG DAN-LK4m-68mpb_92C1R assembled by PacBio and the equivalent MAG assembled from Illumina data only (sharing 99.91 ANI).

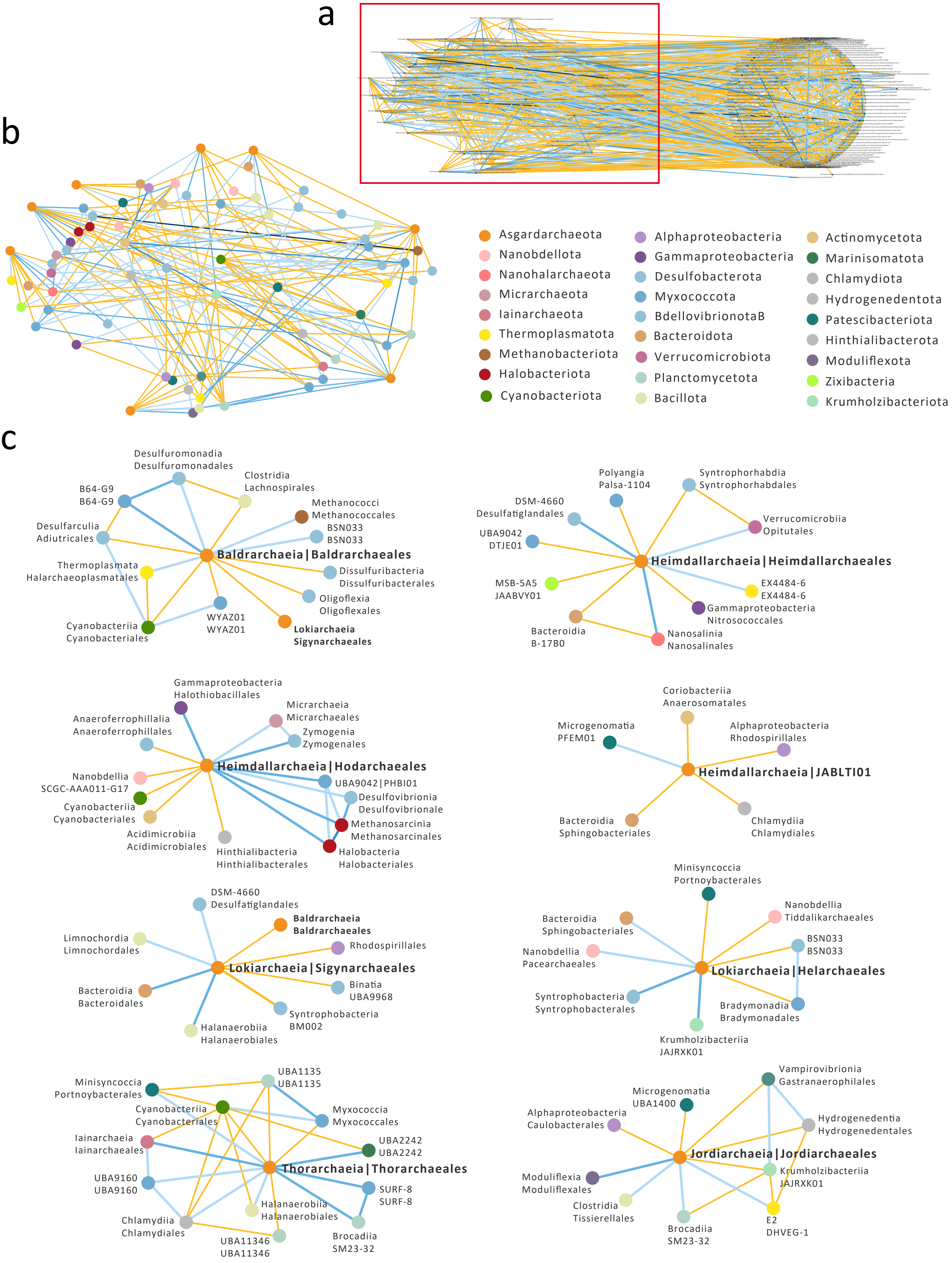


**Supplementary Fig. 5. Co-occurrence network inferred with SparCC based on metagenome-derived USCGs in DAN-LK4 microbial mats.** a, Full co-occurrence network. b, Sub-network (red-square in a) containing Asgardarchaeota nodes and their closest neighbors. c, Ego-networks of Asgardarchaeota grouped at the order level. The class and order of correlating taxa is indicated. Nodes (taxa) are colored according to the phylum they belong to. Edges indicatepositive (blue) or negative (orange) correlations.

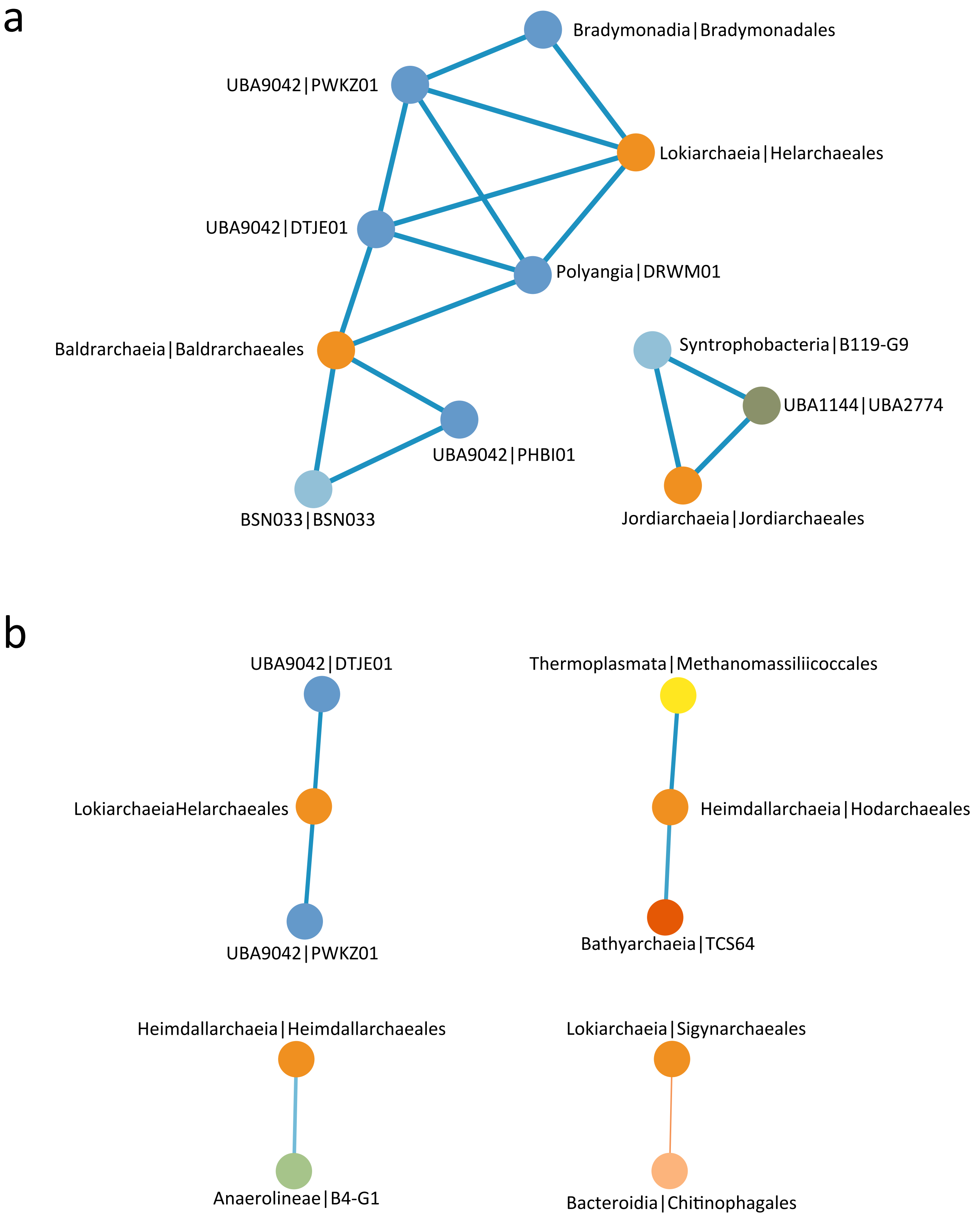


**Supplementary Fig. 6. Co-occurrence network inferred Spearman correlations (a) and SPIEC-EASI based on metagenome-derived USCGs in DAN-LK4 microbial mats.** Only ego-networks of Asgardarchaeota grouped at the order level are shown. The class and order of correlating taxa is indicated. Nodes (taxa) are colored according to the phylum they belong to (color code, same as Supplementary Fig.5). Edges indicatepositive (blue) or negative (orange) correlations.

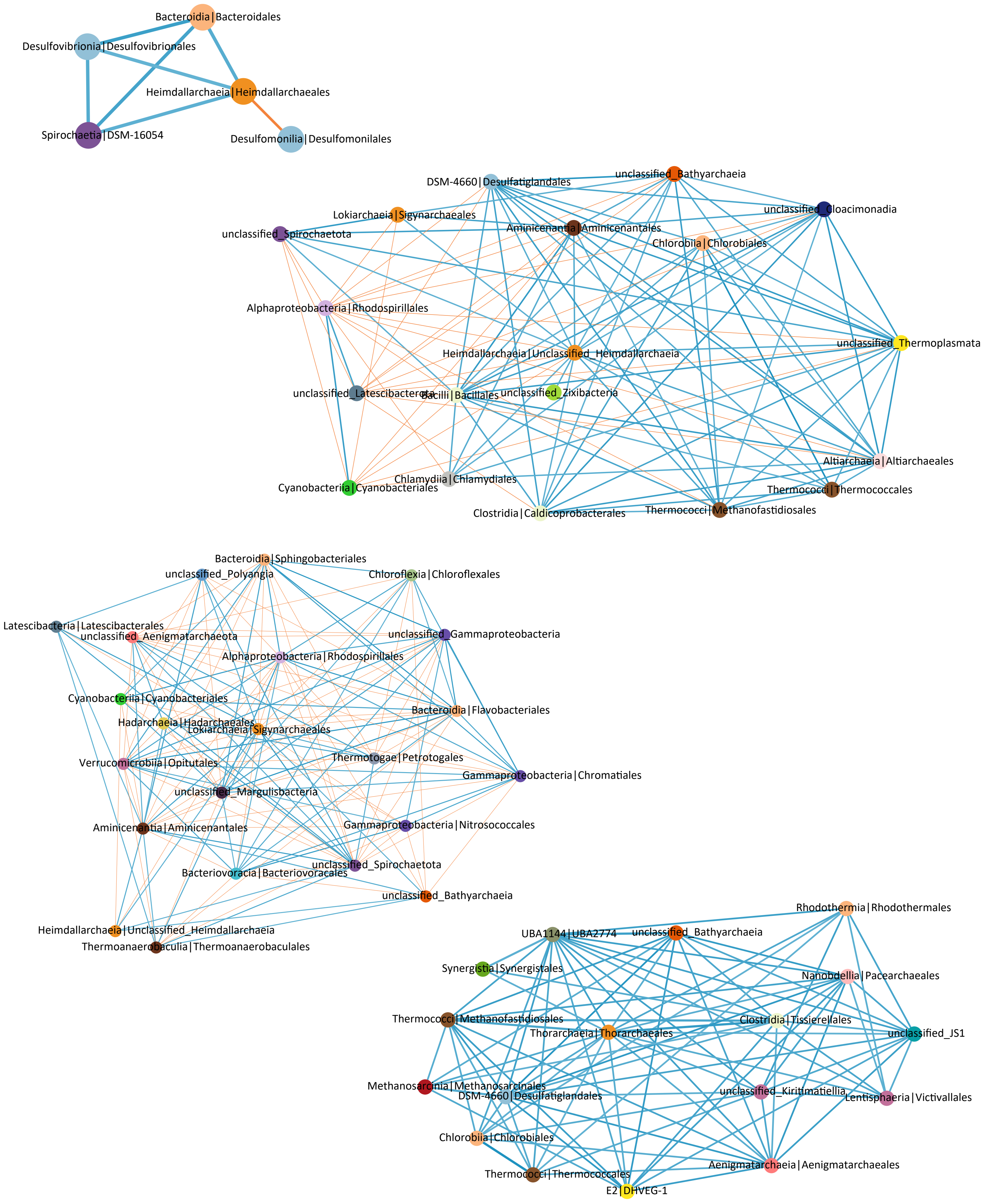


**Supplementary Fig. 7. Co-occurrence network of 16S rRNA gene ASVs in DAN-LK4 microbial mats inferred using Spearman correlations.** Only ego-networks of Asgardarchaeota grouped at the order level are shown. The class and order of correlating taxa is indicated. Nodes (taxa) are colored according to the phylum they belong to (color code as in Supplementary Fig. 5). Edges indicate positive (blue) or negative (orange) correlations.

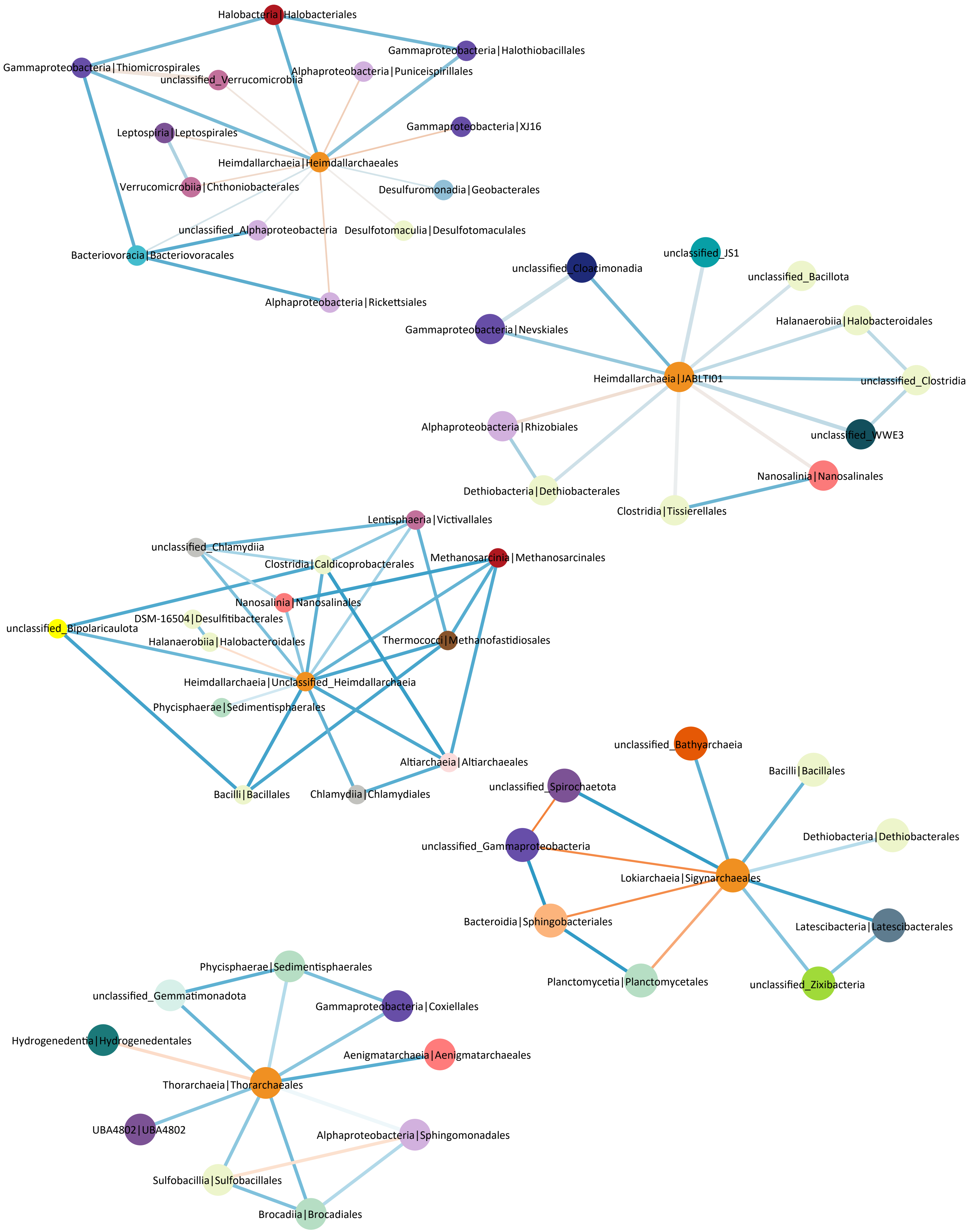


**Supplementary Fig. 8. Co-occurrence network of 16S rRNA gene ASVs in DAN-LK4 microbrial mats inferred using SPIEC-EASI.** Only ego-networks of Asgardarchaeota grouped at the order level are shown. The class and order of correlating taxa is indicated. Nodes (taxa) are colored according to the phylum they belong to (color code as in Supplementary Fig. 5). Edges indicate positive (blue) or negative (orange) correlations.

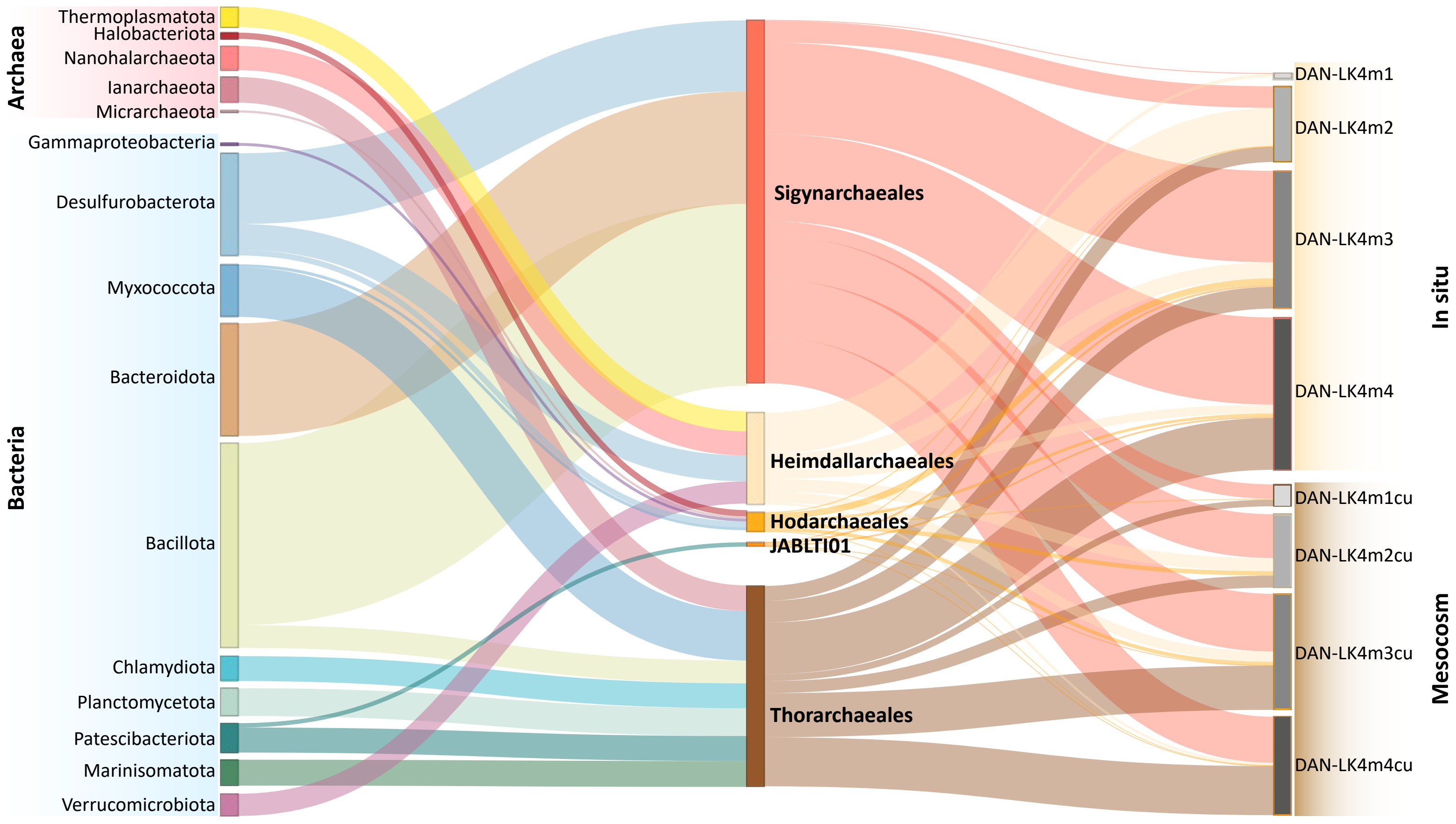


**Supplementary Figure 9. Distribution of asgardarchaeal MAGs assembled from DAN-LK4 microbial mats grouped at order level and their co-occurring taxa along vertical microbial mat strata.** Only the size of asgardarchaeal boxes (center) is proportional to the number of links with prokaryotic taxa (left) and different mat layers (right). The size of boxes of archaea and bacteria on the left are adjusted to the size of the central boxes. Only positive correlations are shown.

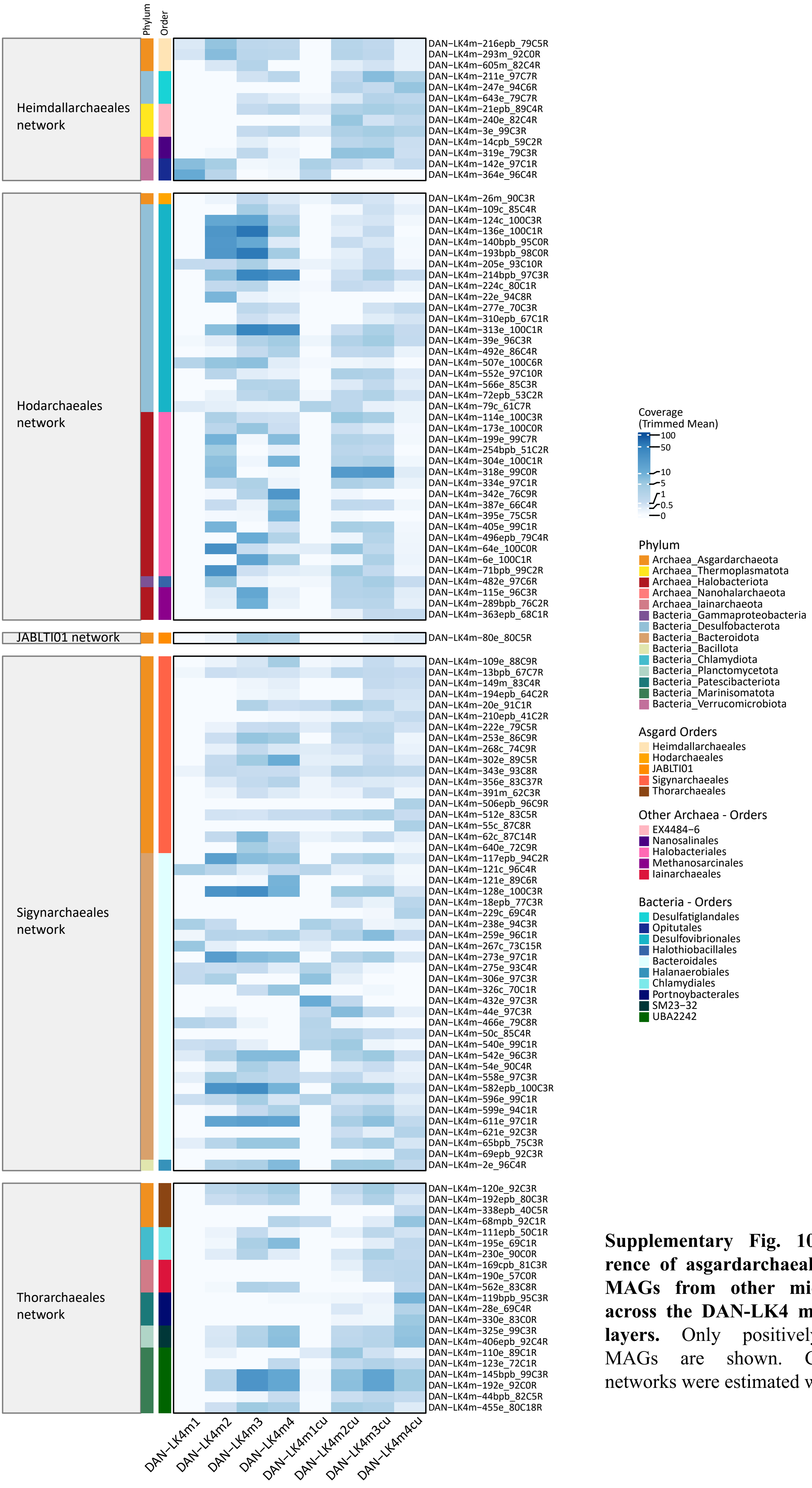


**Supplementary Fig. 10. Co-occurrence of asgardarchaeal MAGs and MAGs from other microbial taxa across the DAN-LK4 microbial mat layers.** Only positively correlated MAGs are shown. Co-occurrence networks were estimated with SparCC.

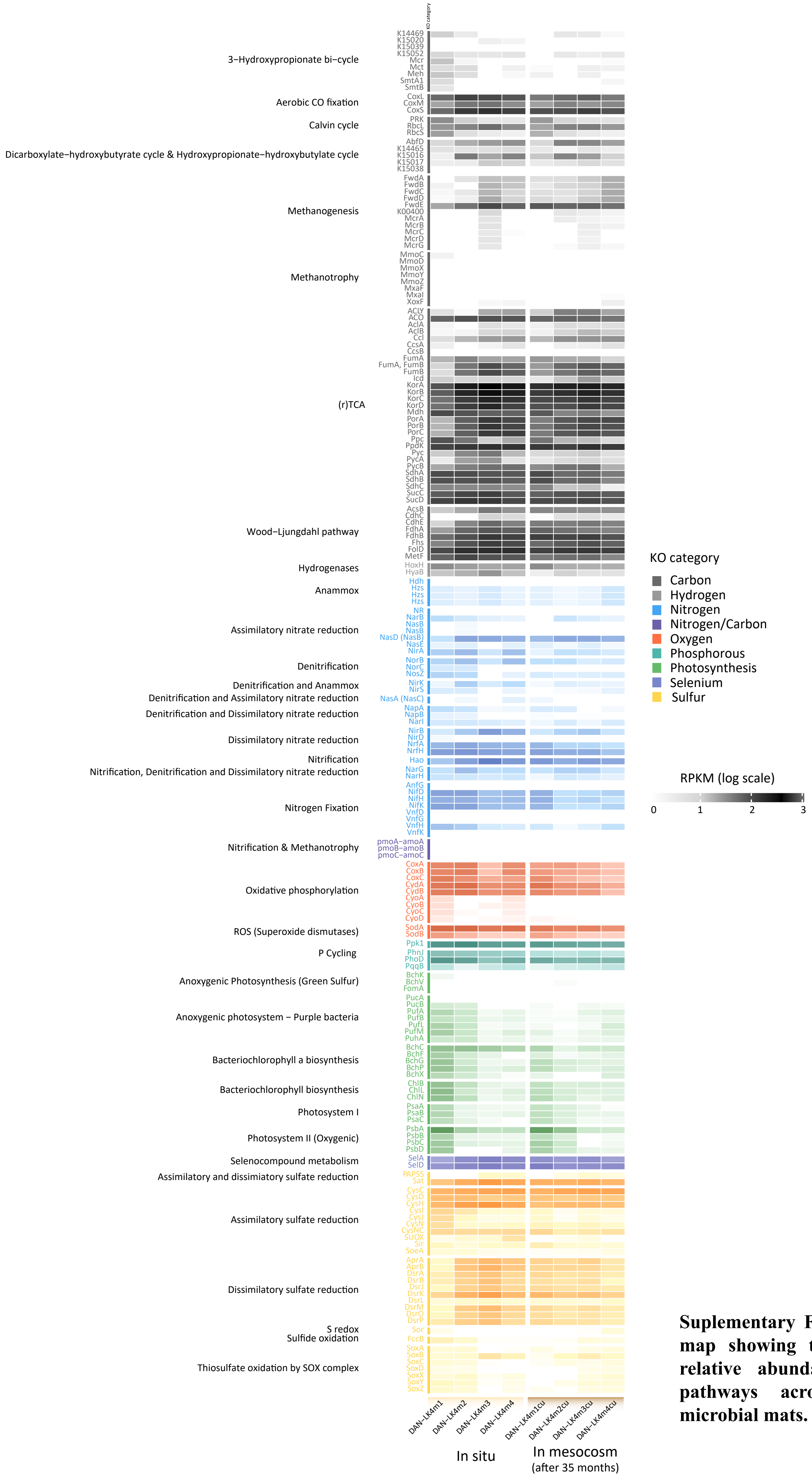


**Suplementary Fig. 11. Detailed heat map showing the distribution and relative abundance of genes and pathways across the DAN-LK4 microbial mats.**

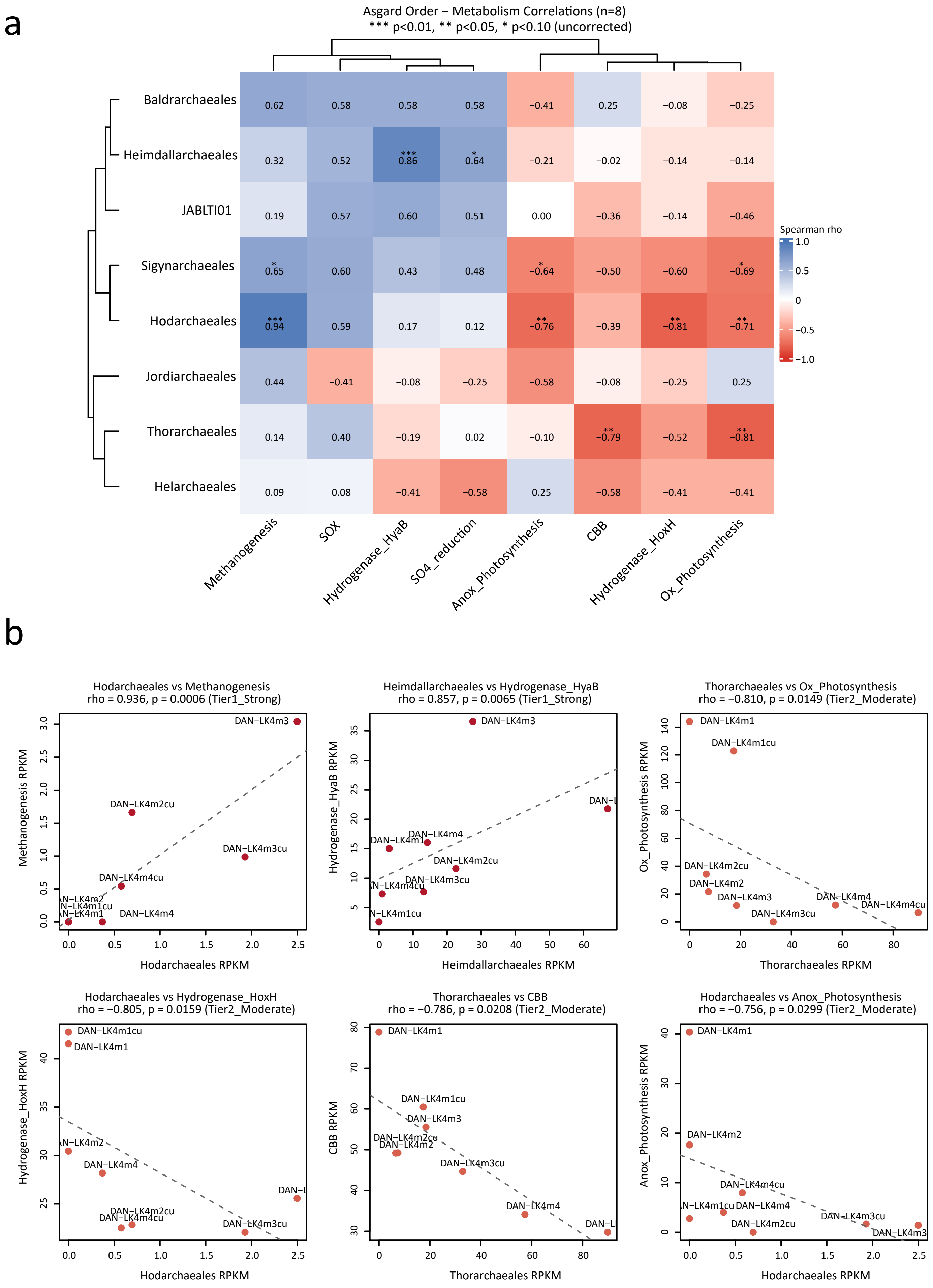


**Supplementary Fig. 12. Statistical correlations of Asgard archaeal orders with major metabolic pathways/genes across the DAN-LK4 metagenomes. a**, Heatmap showing the detailed p-values for Fig.4b. **b**, Pairwise statistical analyses of metabolic genes/pathways and individual asgardarchaeal orders.
